# Identification of chloroplast envelope proteins with critical importance for cold acclimation

**DOI:** 10.1101/813725

**Authors:** Oliver Trentmann, Timo Mühlhaus, David Zimmer, Frederik Sommer, Michael Schroda, Ilka Haferkamp, Isabel Keller, Benjamin Pommerrenig, H. Ekkehard Neuhaus

**Affiliations:** Technische Universität Kaiserslautern, Department of Biology, Plant Physiology, P.O. Box 3049. 67653 Kaiserslautern, Germany; Technische Universität Kaiserslautern, Department of Biology, Computational Systems Biology, P.O. Box 3049, 67653 Kaiserslautern, Germany; Technische Universität Kaiserslautern, Department of Biology, Molecular Biotechnology & Systems Biology, 67653 Kaiserslautern, Germany

## Abstract

The ability of plants to cope with cold temperatures relies on their photosynthetic activity. This already demonstrates that the chloroplast is of utmost importance for cold acclimation and acquisition of freezing tolerance. During cold acclimation, the properties of the chloroplast change markedly. To provide the most comprehensive view of the protein repertoire of chloroplast envelope, we analysed this membrane system in *Arabidopsis thaliana* using MS-based proteomics. Profiling chloroplast envelope membranes was achieved by a cross comparison of protein intensities across plastid and the enriched membrane fraction both under normal and cold conditions. Multivariable logistic regression models the probabilities for the classification problem to address envelop localization. In total, we identified 38 envelope membrane intrinsic or associated proteins exhibiting altered abundance after cold acclimation. These proteins comprise several solute carries, such as the ATP/ADP antiporter NTT2 (substantially increased abundance) or the maltose exporter MEX1 (substantially decreased abundance). Remarkably, analysis of the frost recovery of *ntt* loss-of-function and *mex1* overexpressor mutants confirmed that the comparative proteome is well suited to identify novel key factors involved in cold acclimation and acquisition of freezing tolerance. Moreover, for proteins with known physiological function we propose scenarios explaining their possible role in cold acclimation. Furthermore, spatial proteomics introduces a novel layer of complexity and enabled the identification of proteins differentially localized at the envelope membrane under the changing environmental regime.

## Introduction

No other plant cell organelle is so typically associated to the autotrophic lifestyle than the chloroplast. Chloroplasts of higher plants evolved from cyanobacteria and still contain a small, endogenous genome mainly encoding proteins located to the thylakoid membrane system (McFadden, 1999). These organelles are the cellular site of photosynthetic light reactions and oxygen release, they harbour the enzymatic machineries required for photosynthetic CO_2_ fixation, for starch production, nitrite and sulphate reduction, and amino acid- and fatty acid synthesis (Buchanan, 2015).

To fulfil all these functions chloroplasts, must import and export a wide variety of metabolic intermediates (Weber et al., 2005) and they have to communicate with the nucleus to balance plastidic and nuclear gene expression (Pfalz et al., 2012). Accordingly, changing external parameters, such as light intensities or temperatures, result in substantial genetic and metabolic re-adjustments and it has been proposed that chloroplasts even serve as “sensors”, centrally positioned in the plants reaction to abiotic stress stimuli (Crosatti et al., 2013). The required molecular communication between chloroplasts and the nucleus takes place via retrograde and anterograde signalling processes (Kleine and Leister, 2013), while the altered metabolite exchange between the chloroplast and the cytosol depends among others upon corresponding changes in the envelope proteome.

Cold tolerant species can gain the capacity to survive freezing temperatures by a process termed “cold acclimation” (Catalá et al., 2011), starting when plants face cold but non-freezing temperatures. Accordingly, *Arabidopsis thaliana*, as a typical cold hardy species (Yano et al., 2005), represents a suitable tool for the investigation of molecular and physiological mechanisms underlying acclimation to low temperatures (Strand et al., 1997; Calixto et al., 2018; Nägele and Heyer, 2013; Schulze et al., 2012; Rekarte-Cowie et al., 2008).

For following considerations, it appears likely that changes in the chloroplast envelope contribute to the ability of higher plants to acclimate rapidly to decreasing external temperatures. First, proper acclimation to cold depends on photosynthetic activity, providing sugars required for the toleration of low temperatures (Wanner and Junttila, 1999; Alberdi and Corcuera, 1991; Pommerrenig et al., 2018). Therefore, under cold conditions, chloroplasts must maintain, although at lower rates, the daytime export of triose-phosphates to allow sugar synthesis in the cytosol. Second, nocturnal starch degradation leads to the presence of glucose and maltose in the chloroplast stroma (Kötting et al., 2010; Sicher, 2011) and mutants with impaired starch mobilization exhibit less freezing tolerance (Kaplan and Guy, 2004; Yano et al., 2005). Thus, we suppose that after onset of chilling temperatures, the export of glucose and maltose must be adapted to altered starch turnover. Third, for effective cryoprotection of thylakoid membranes raffinose must be imported into the chloroplast (Schneider and Keller, 2009; Knaupp et al., 2011). Fourth, in the cold, stromal sucrose is relocated to the cytosol where it contributes to the acquisition of a maximal freezing tolerance (Patzke et al., 2019). Fifth, to maintain enough membrane fluidity at low temperatures, cold acclimation induces the remodelling of structural lipids in thylakoids and envelope membranes (Barrero-Sicilia et al., 2017; Moellering et al., 2010).

To investigate putative changes in the protein composition caused by exposure of plants to cold temperatures several proteomic analyses have been carried out using in most cases total leaf extracts from the model plant Arabidopsis, the closely related species Thellungiella and also crop plants like Alfalfa and Wheat (Gao et al. 2009; Rocco et al. 2013; Amme et al. 2006; Awai et al. 2006; Chen et al. 2015; Kosová et al. 2013). Furthermore, the lumen and stromal proteome of isolated Arabidopsis chloroplast has been examined after plant exposure to 5°C for different time periods (Goulas et al. 2006).

Although our knowledge on cold-induced metabolic changes in chloroplasts and associated processes is quite comprehensive, it is completely unknown whether and to which degree alterations in the abundance of envelope located proteins contribute to cold acclimation. During the past two decades, the protein composition of the envelope membrane has been examined intensively. Particularly, studies by Ferro and co-workers (Ferro et al., 2003; Ferro et al., 2010) supported the establishment of AT_CHLORO, a comprehensive and experimentally substantiated open access database for sub-plastidic protein localization (Bruley et al., 2012). This work has been extended very recently by a study unrevealing so far hidden components of the envelope membrane (Bouchnak et al. 2019)

To our knowledge, comparative studies reporting on alterations in the protein composition of the envelope membrane caused by environmental changes are missing. Because of the central role of the chloroplast in cold acclimation, we hypothesized that during this process, the protein content and composition of the envelope becomes modified. Therefore, we performed a comparative analysis of envelope proteins from cold treated plants and from plants permanently grown under standard conditions. We decided to conduct a label-free quantitative proteome study because the labelling of proteins during plant growth (e.g. ^15^N labelling) requires hydroponic cultivation whereas plants for the label-free proteome study can be grown on soil and thus under more natural conditions. It is important to mention that the label-free approach has its specific limitations. However, these limitations can be minimized using more biological replicates and extensive statistical analyses (Trentmann and Haferkamp, 2013).

To affirm the physiological relevance of identified changes in protein abundances we exemplarily investigated the gain of frost tolerance after cold acclimation for two candidate proteins with opposite abundance changes using loss of function or gain of function mutants.

## Results

### Envelope membrane purification and mass spectrometry

The isolation of intact chloroplasts from cold acclimated plants and control plants was performed according to Kunst et al. (Kunst, 1998) with some marginal modifications. Particularly, the initial disruption of the leave tissue with a commercial electric blender turned out to be a critical step since even slightly prolonged pulsing periods resulted in a dramatically decreased chloroplast yield. The intactness of chloroplasts was determined by phospho-glucose-isomerase (PGI) enzyme assay. PGI activity of intact was normalized to that of disrupted (set to 100%) chloroplasts. The intactness generally ranged between 85% and 95% (Supp. Fig. S1B). This observation suggests that the quality of the isolated chloroplasts is not affected by the used cultivation temperatures (see also Supp. Fig. S1A), an important prerequisite for the subsequent isolation of the envelope membranes. The three-step sucrose gradient led to a yellowish band without visible chlorophyll contaminations, which is indicative for no, or only minor contaminations with thylakoid membranes. The appearance of this envelope fraction generally resembled that reported by Ferro (Ferro et al., 2003). Because of the increased detection sensitivity of mass spectrometry, we could reduce the amount of leaf material per single isolation from about 500 g (as reported by Ferro et al. 2003) to 200 g. Purified envelope membranes were collected by ultracentrifugation and finally washed five-times with 1M sodium carbonate. Sodium carbonate treatment allows removal of soluble proteins weakly attached to membranes (Kim et al., 2015). By this approach, we obtained about 5 µg envelope membrane proteins from 200 g Arabidopsis leaves. Proteins of total chloroplast lysates and of the envelope membrane fraction from cold-acclimated and non-acclimated plants were separated by SDS-Page. Subsequent to in-gel tryptic digestion, the resulting peptides in the different samples were analysed by nanoLC-MS/MS. The proteome analysis of chloroplasts and envelopes from cold and non-acclimated plant identified 905 proteins in total (Supplement Table S1).

**Table 1:** Intrinsic or chloroplast envelope associated proteins with changed abundances after 4 days of cold acclimation at 4°C. **TM**: number of transmembrane domains, revised using information provided by ARAMEMNON release 8.1 and protein specific publications; **Local.**: localisation of the identified proteins based on the AT_CHLORO and this study, **Ch**: chloroplast, **E**: envelope, **IE**: inner envelope, **IO**: outer envelope, **S**: stroma **\***: previously predicted envelope localisation confirmed by this study, **new**: by this study identified intrinsic or envelope associated proteins

| Gene ID | Protein names |  | log2FC<br>4°C - 22°C | Stdv.<br>log2FC<br>change | TM | Local. |
| --- | --- | --- | --- | --- | --- | --- |
| *At1g15500* | plastidic ATP/ADP antiporter (AATP2/NTT2) | ↑ | +9.69 | 0.33 | 11 | Ch/E/IE |
| *At4g17170* | putative RAB-B-class small GTPase (RAB-B1b) | ↑ | +7.46 | 0.19 | 0 | Ch/E new |
| *At5g64840* | putative subfamily F ABC protein (ABCF5/GCN5) | ↑ | +10.07 | 0.38 | 0 | Ch/E new |
| *At5g33320* | phosphoenolpyruvate/phosphate translocator (PPT1/CUE1) | ↓ | -11.06 | 0.45 | 7-8 | Ch/E/IE |
| *At5g17520* | maltose translocator (RCP1/MEX1) | ↓ | -9.05 | 0.27 | 9 | Ch/E/IE |
| *At3g51140* | putative DnaJ-chaperone-like protein | ↓ | -1.05 | 0.41 | 4 | Ch/E/OE |
| *At4g39460* | S-adenosylmethionine transporter (SAMC1/SAMT1) | ↓ | -2.07 | 0.76 | 6 | Ch/E/IE |
| *At5g16010* | putative steroid 5-alpha reductase | ↓ | -10.18 | 0.12 | 6-7 | Ch/E * |
| *At4g32400* | nucleotide uniporter (SHS1/BT1) | ↓ | -1.10 | 0.44 | 6 | Ch/E/IE |
| *At5g42960* | putative OEP24-type outer membrane channel | ↓ | -1.14 | 0.47 | 0<br>12 β-barrels | Ch/E/OE |
| *At1g65410* | putative component of ER-to-thylakoid lipid transfer complex (TGD3/ABCI13/NAP11) | ↓ | -9.25 | 0.52 | 0 | Ch/E/IE |
| *At4g33350* | chloroplast inner envelope translocon component (Tic22-IV) | ↓ | -1.23 | 0.27 | 0 | Ch/E/IE |
| *At3g01500* | Beta carbonic anhydrase 1, chloroplastic | ↓ | -1.32 | 0.34 | 0 | Ch/E new |
| *At2g01320* | putative subfamily G ABC-type transporter (ABCG7/WBC7) | ↓ | -9.52 | 0.54 | 5 | Ch/E * |
| ***At5g14220*** | protoporphyrinogen IX oxidase (PPO2) | ↓ | **-1.44** | 0.57 | 0 | Ch/E * |
| ***At5g12860*** | plastidic 2-oxoglutarate/malate translocator (DiT1/pOMT1) | ↓ | **-1.45** | 0.90 | 14 | Ch/E/IE |
| ***At1g08640*** | CJD1 (Chloroplast J-like Domain 1) influences fatty acid composition of chloroplast lipids | ↓ | **-1.47** | 0.37 | 3 | Ch/E/IE |
| ***At5g02940*** | putative Pollux/Castor-type voltage-gated ion channel (Pollux-L1) | ↓ | **-1.48** | 1.20 | 3 | Ch/E * |
| ***At2g43630*** | protein of unknown function | ↓ | **-1.53** | 0.37 | 1 | Ch/E/IE |
| ***At3g08740*** | Elongation factor P (EF-P) family protein | ↓ | **-7.60** | 0.86 | 0 | Ch/E &<br>Ch/S |
| ***At1g78620*** | putative phytyl-phosphate kinase (VTE6) | ↓ | **-1.56** | 0.09 | 6 | Ch/E/IE |
| ***At5g23040*** | putative DnaJ-chaperone-like protein involved in protochlorophyllide oxidoreductase stabilization (CPP1/CDF1/DnaJD11) | ↓ | **-1.58** | 0.11 | 3-4 | Ch/E/IE |
| ***At2g45740*** | member of the peroxin11 (PEX11) gene family | ↓ | **-7.66** | 0.37 | 1-3 | Ch/E/IE |
| ***At1g10510*** | RNI-like superfamily protein EMBRYO DEFECTIVE 2004 | ↓ | **-0.65** | 0.20 | 1 | Ch/E/IE |
| ***At3g10840*** | putative alpha/beta-fold-type hydrolase | ↓ | **-9.74** | 0.36 | 0-2 | Ch/S |
| ***At3g32930*** | protein of unknown function | ↓ | **-7.72** | 0.39 | 0 | Ch/S |
| ***At2g17695*** | putative chloroplast outer envelope solute channel (OEP23) | ↓ | **-7.74** | 0.44 | 0 | Ch/E * |
| ***At2g42770*** | putative PMP22/Mpv17-type protein of unknown function | ↓ | **-0.71** | 0.09 | 2-4 | Ch/E/IE |
| ***At3g20330*** | aspartate carbamoyltransferase (ATCase) | ↓ | **-9.80** | 0.87 | 0 | Ch/E/IE |
| ***At2g24820*** | putative component of inner envelope protein import machinery \|<br>phylobilin hydroxylase (TIC55/Tic55-II) | ↓ | **-0.76** | 0.37 | 2 | Ch/E/IE |
| ***At3g57280*** | plastid fatty acid exporter (FAX1) | ↓ | **-1.77** | 0.31 | 4 | Ch/E/IE |
| ***At5g42130*** | putative (animal Mitoferrin)-like carrier (MFL1) | ↓ | **-0.79** | 0.20 | 6 | Ch/E/IE |
| ***At3g49560*** | putative tRNA import component of mitochondrial membrane translocase machinery (TRIC1/PRAT2.1/HP30-1) | ↓ | **-0.81** | 0.27 | 2-4 | Ch/E/IE |
| *At3g56910* | putative plastid-specific ribosomal protein (PSRP5) | ↓ | **-0.85** | 0.59 | 0 | Ch/E/IE |
| *At4g28620* | putative subfamily B ABC-type transporter (ABCB24/ATM2) | ↓ | **-0.88** | 0.73 | 6 | Ch/E new |
| *At3g63410* | methyl-6-phytyl-1,4-hydroquinone methyltransferase (APG1/VTE3) | ↓ | **-0.92** | 0.62 | 1 | Ch/E/IE |
| *At3g20320* | putative substrate-binding component of ER-to-thylakoid lipid transfer complex (TGD2/ABCI15) | ↓ | **-1.99** | 1.71 | 1 | Ch/E/IE |
| *At2g35800* | putative calcium-dependent S-adenosyl methionine carrier (SAMTL) | ↓ | **-0.99** | 0.39 | 2 | Ch/E * |

### Envelope membrane protein profiling

We used spatial proteomics for envelope membrane protein profiling. The principle of this technique is to fractionate organelles or sub-compartments and to identify the distribution of proteins across the differentially enriched sub fractions. Here, we compared the protein occurrence in the chloroplast fraction with that in the envelope fractions from standard and cold cultivated plants. Envelope located proteins generally should be present in both fractions (total chloroplast lysate and enriched envelopes) but should be enriched in the envelope preparation, whereas non-envelope proteins should be depleted from this fraction. Consequently, comparing protein abundances in the two fractions already allows the calculation of an enrichment factor for the envelope located candidates. In order to model the probabilities for the classification of the identified proteins, either envelope or not, we applied multivariable logistic regression. Combined information from AT_CHLORO (Bruley et al., 2012) and the Plant Proteome Database (Sun et al., 2009) revealed high quality localization data for 453 of the 905 identified proteins, with 162 being assigned to the envelope and 291 to an alternative location (either stroma or thylakoid). The corresponding proteins were chosen as markers for the training of our classifier (localization data and enrichment factors).

Cold temperatures are known to alter the lipid content and composition of cellular membranes in plants. In order to exploit possible temperature-induced changes in the membrane structure affecting protein extraction, we used both enrichment factors and additionally incorporated a subset of physiochemical amino-acid properties with minimal redundancy, while retaining maximum relevance using a minimum spanning tree approach (Zimmer et al., 2018). After training, we reached 97.82 % prediction accuracy accessed by 10-fold cross validation.

The graphical representation of the score distribution demonstrates that the chosen parameters allows the discrimination between envelope and non-envelope located proteins with quite high accuracy (Figure 1). Most non-envelope proteins exhibit negative score values whereas envelope proteins occur with higher frequency at positive scores. Consequently, proteins with ambiguous or unknown localization can be assigned to the envelope or non-envelope group according to their individual score values. We analysed the score behaviour of all 905 identified proteins to check for eventual envelope localization. For this, we tolerated a false discovery rate of 5% and thus the score value of 0.92 was chosen as cut-off. By this strategy, 207 of the originally identified proteins could be annotated as envelope located (Supplement Table S1 and Figure 2). The overall quality of our proteome study is given by the pie chart in Figure 2. Here the localization of the identified proteins described by hierarchically organized localization ontology.

**Figure 1:**
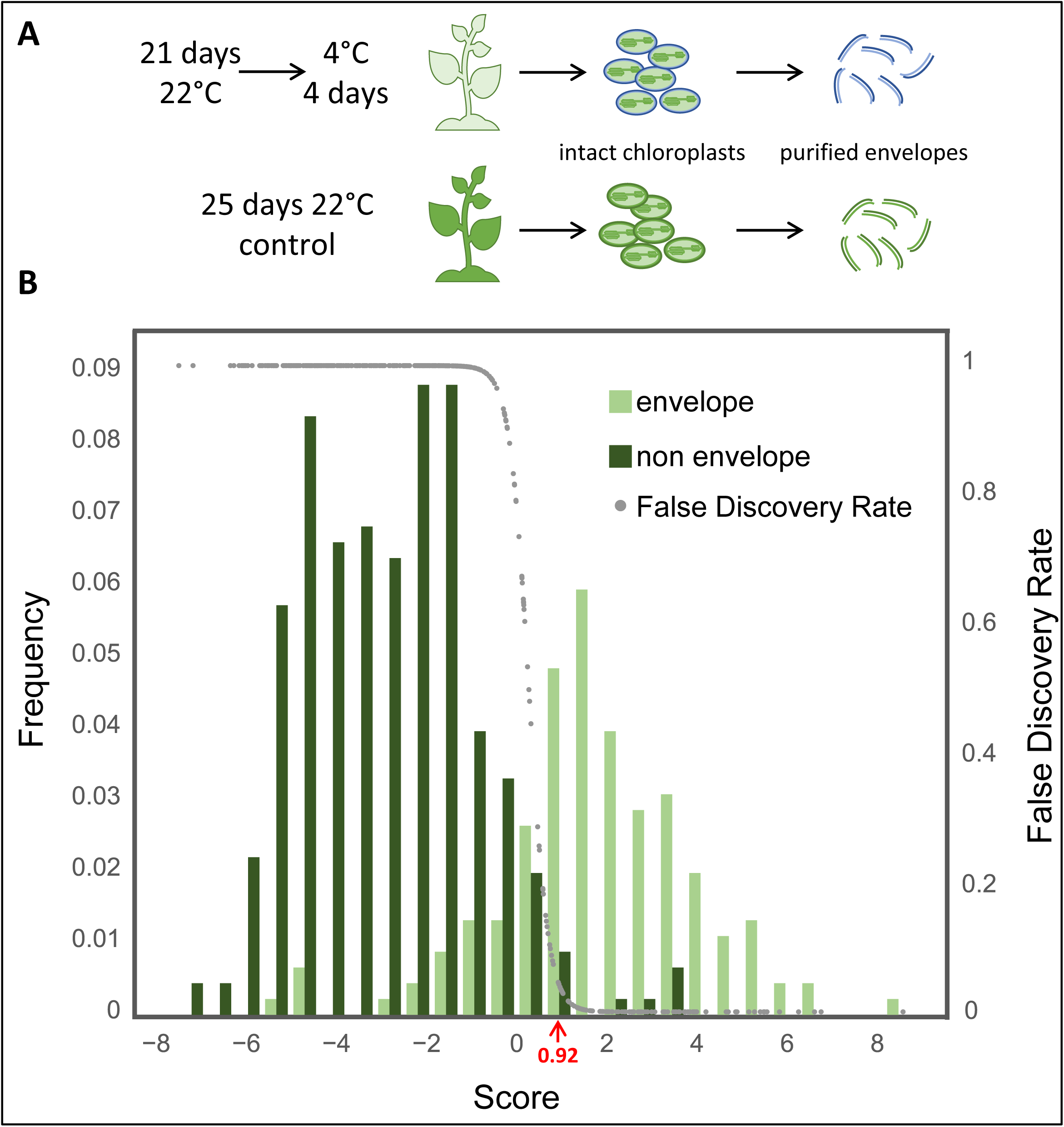
A. Schematic drawing illustration cold acclimation and organelle/envelope isolation. **B** Graphical representation of the score distribution learned from protein features with known envelope localization (light green) and proteins from other compartments (dark green). The divergence between the distributions allows separating the two different classes by the score behavior based on the selected protein features. False discovery rate (grey) was calculated after 10-fold cross-validation using the knowledge from all previously known protein localization in the data set.

**Figure 2:**
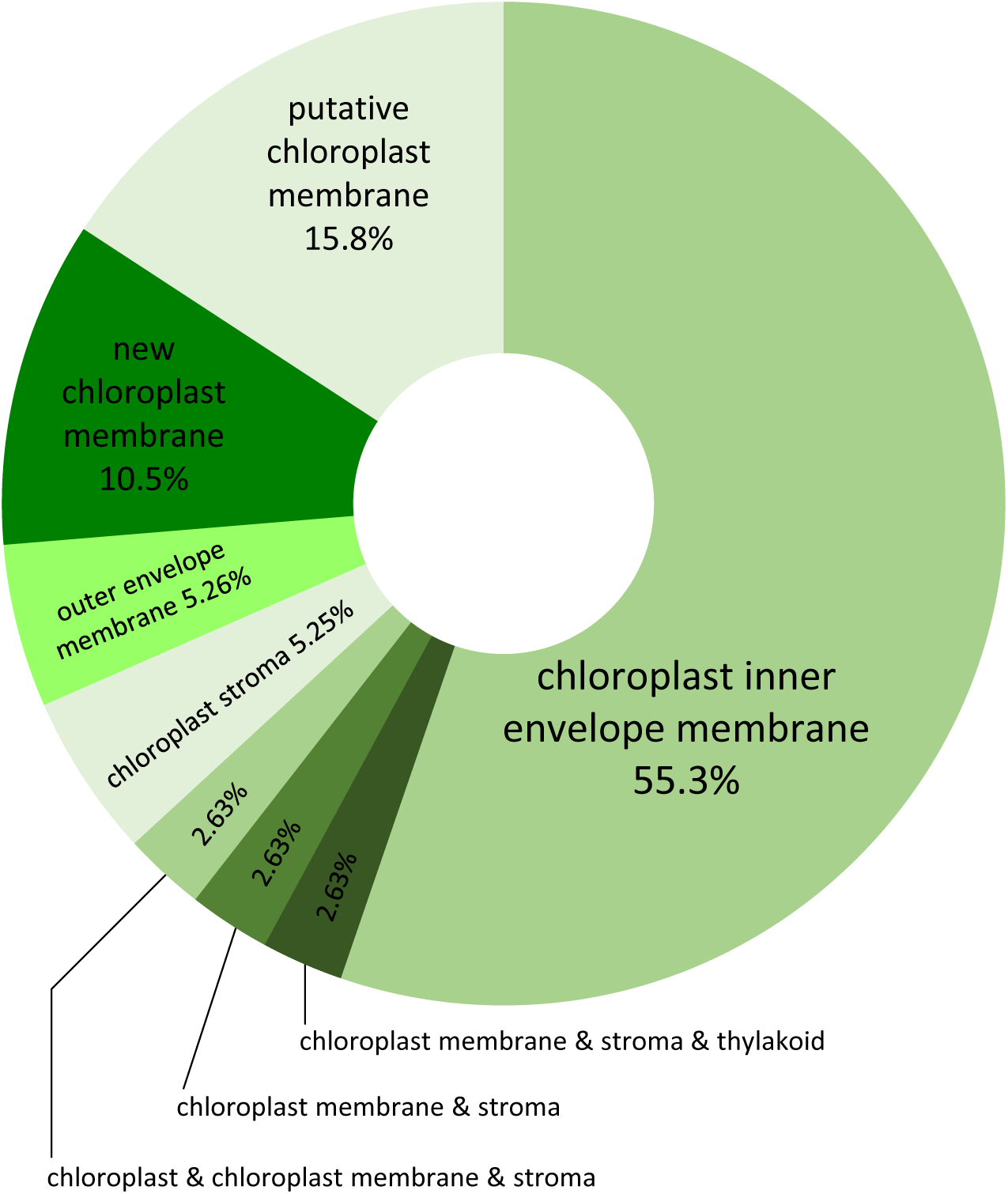
Pie chart demonstrating the general quality of the performed envelope proteome study. Localization of the identified proteins described by hierarchically organized localization ontology. Multiple ontology tags indicate either technical indistinguishable or a biological meaningful multi-localization of the respective proteins.

### Cold temperatures alter the protein repertoire of the envelope membrane

The central role of the chloroplast in cold acclimation led us to the assumption that exposure of Arabidopsis to low temperatures might alter the amount and composition of proteins in the envelope membrane. Accordingly, envelopes from cold treated plants might contain proteins that have not been assigned to this localization before, because they are missing or of very low abundance and thus were not detected in the chloroplast envelope of plants grown under optimal culture conditions. Moreover, one might envision that certain chloroplast proteins are differently located under varying environmental regimes. Spatial proteomics allow insights into the cellular organization as well as in the dynamics of the subcellular distribution of proteins and might also help to identify novel envelope proteins, particularly in the cold treated plants.

First, we compared the envelope protein levels of cold acclimated plants with those of control plants. This analysis revealed that cold treatment changed the abundance of approximately 20% (38 of 207) of the identified envelope proteins (Table 1). Most of these proteins (35 out of 38) showed lower abundance. It is imaginable that cultivation of plants under cold temperatures decreases the amount of the detected envelope proteins either due to a generally decelerated protein synthesis or due to lesser efficient extraction from the corresponding membranes. However, the facts that the individual proteins exhibit different degrees in reduction and that at least three proteins were even of substantially higher abundance (log2FC from 7.46 to 10.07) contradicts this assumption (Table 1). Interestingly, one of the three proteins with increased abundance was previously not assigned to the envelope location (ABCF5/GCN5, *At5g64840*). Moreover, also the set of proteins that decreased after cold treatment contained two novel envelope candidates (BCA1, *At3g01500*; ABCB24/ATM2, *At4g28620*).

To analyse whether the differently abundant envelope proteins are intrinsic to the membrane we checked for possible membrane spanning domains. For this, we manually curated the information from AT_Chloro, and the Plant Proteome Database with data obtained from ARAMEMNON (release 8.1) and diverse publications. For instance, the list of envelope membrane proteins (Table 1) contains four members of the mitochondrial carrier family (BT1 *At4g32400*; SAMC1, *At4g39460*; SAMTL, *At2g35800* and MFL1, *At5g42130*). These carriers were initially proposed to contain no or a lower number of TMs. However, members of this protein family are classified by their common basic structure, which amongst others comprises six transmembrane domains (TMs) (Haferkamp and Schmitz-Esser, 2012). Therefore, we corrected the corresponding information accordingly. Manual curation of the previous data led to the identification of alpha helical domains in 26 out of the 38 envelope proteins. The list of differentially abundant proteins also contains two outer envelope proteins (OEP23 *At2g17695* and OEP24 *At5g42960*). OEP23 and OEP24 exhibit amphiphilic helices or a ß-barrel confirmation and act as cation and anion channels, respectively (Goetze et al., 2015; Röhl et al., 1999). Accordingly, at least 28 of the 38 differentially abundant envelope proteins show features of membrane intrinsic proteins.

The remaining ten proteins are considered as rather soluble and the fact that they were not removed by sodium carbonate suggests that they are tightly attached to the membrane. This membrane association might result from a specific interaction with a membrane intrinsic protein. TGD3 (*At1g65410*) for example represents the ATPase subunit of the lipid transporter TGD and by this is apparently fixed to the membrane. Moreover, Tic22-IV is part of the protein import machinery, and thus might interact not only physiologically but also physically with membrane components of the TIC complex. RAB-B1b is a putative RAB-B-class small GTPase. Generally, RAB proteins are post-translationally modified by prenylation and RAB-B1 contains a geranylgeranylation motif (Maurer-Stroh and Eisenhaber, 2005). Consequently, RAB-B1, just like other RAB proteins, can be considered as a peripheral membrane protein, which is temporarily anchored to a membrane via its lipid group but can be released from this location during the GTPase cycle. Interestingly, a proteome study revealed that the putative aspartate carbamoyltransferase is palmitoylated (Hemsley et al., 2013) and thus might be attached to the envelope via its lipid anchor. Moreover, a palmitoylation site is predicted for the putative plastid-specific ribosomal protein (PSRP5 *At3g56910*) (Ren et al., 2008). Since the remaining five proteins lack clearly predicted lipid modification motifs, their membrane association might be caused by an interaction with a membrane protein.

By the help of the known or predicted physiological function, we aimed to affiliate the differently abundant envelope proteins to functional groups. A high number (19 out of the 38) of the envelope proteins are associated to the metabolite and protein translocation. The substrates of the corresponding proteins are heterogeneous and range from ions (like potassium of Pollux-L1; *At5g02940*) to comparatively large and complex molecules, like lipids (TGD subunits TGD2 and 3) or protein precursors. From the 19 transport associated envelope proteins only ATP/ADP transporter NTT2 (*At1g15500*; log2FC +9.7) increased whereas the majority decreased after onset of cold (Table 1). The lipid transporter subunit TGD3, the ABC-type transporter ABCG7 (*At2g01320*), OEP23, the maltose exporter MEX1 (*At5g17520*) and the phosphoenolpyruvate/Pi exchanger PPT1 (*At5g33320*) are substantially decreased in abundance (log2FC −7.74 to - 11.06). By contrast, the remaining envelope proteins associated to translocation rather exhibit minor reductions in their abundance, ranging from log2FC of −0.76 for the phyllobilin hydroxylase TIC55-II (*At2g24820*), a putative component of the TIC machinery to log2FC of −2.07 for the S-adenosylmethionine transporter SAMC1 (*At4g39460*).

Interestingly, even though both, NTT2 (*At1g15500*) and BT1 (*At4g32400*), accept adenine nucleotides as substrates cold exposure led to opposed changes in their abundances (Table 1). In this context, however it is important to mention that they fulfil different physiological functions. NTT2 imports ATP in exchange with ADP plus phosphate and by this provides chemical energy to the plastid (Tjaden et al., 1998; Kampfenkel et al., 1995; Reinhold et al., 2007; Trentmann et al., 2008) whereas BT1 represents an uniporter and exports newly generated adenine nucleotides to the cytosol (Kirchberger et al., 2008).

The chloroplast inner envelope harbours three sugar transport proteins, the glucose transporter pGlcT (*At5g16150*), the sucrose exporter pSuT (*At5g59250*) and the maltose exporter MEX1 (*At5g17520*) (Weber et al., 2000; Patzke et al., 2019; Niittylä et al., 2004). Although sugars play an important role in cold acclimation (Pommerrenig et al., 2018; Kaplan et al., 2006) and although all three sugar transporters have been identified in the envelope proteome (Supplement Table S1), solely the abundance of MEX1 changed and in fact became reduced by the cold treatment (Table 1; log2FC −9.0).

Out of the 38 differentially abundant proteins four are clearly involved in fatty acid and lipid metabolism (Table 1). The J-like protein CJD1 (*At1g08640*) influences the composition of chloroplast lipids (Ajjawi et al., 2011) whereas TGD2 (*At3g20230*) and TGD3 (*At1g65410*) represent subunits of the phosphatidic acid transfer complex TGD (Lu et al., 2007), and FAX1 (*At3g57280*) mediates fatty acid export (Li et al., 2015a). Moreover, because the α/β hydrolase superfamily comprises proteases, dehalogenase, peroxidases as well as epoxide hydrolases, lipases and esterases, the putative alpha/beta-fold-type hydroalase (*At3g10840*) might also be associated to lipid and sterol metabolism (Ollis et al. 1992). While the latter enzyme and TGD3 are highly reduced in their abundance, the remaining three proteins showed comparatively low decrease during cold exposure. Moreover, also the cytosolic located RAB-B-class small GTPase RAB-B1b (*At4g17170*) might indirectly join the group of lipid metabolism associated proteins, since this class of proteins modifies intracellular membrane fluxes and by this lipid composition (Karim and Aronsson, 2014). The alterations in the abundances of envelope proteins involved in lipid homeostasis might be causative for cold-induced changes in the membrane lipid composition of the chloroplast and the surrounding cell (Barrero-Sicilia et al., 2017).

Tocopherols are cellular antioxidants that protect fatty acids from peroxidation and by this may stabilize chloroplast membranes also during freezing (Hincha, 2008). Therefore, it was surprising that two proteins of the tocopherol biosynthesis, VTE6 (*At1g78620*) and VTE3 (*At3g63410*) were of lower abundance in cold treated plants (Mène-Saffrané, 2017; Fritsche et al., 2017). Apart from one enzymatic reaction, the synthesis of vitamine E components (comprising tocopherols, tocotrienols and plastochromanols) takes place at the inner envelope membrane (Cheng et al., 2003; van Wijk and Kessler, 2017). Because of their role in the protection of fatty acids, VTE3 and VTE6 were affiliated to the functional group of envelope proteins associated to membrane lipid modification.

Finally, 24 proteins showed a quite moderate alteration in the abundance (log2FC between −2 and +2) whereas 14 show substantial changes (log2FC < −7 or > 7) and include all three proteins that increase in response to cold. To investigate whether the obtained data allow insights in the physiological relevance of altered proteins in cold acclimation we analysed two proteins with opposed changes in their abundance in more detail.

### Cold acclimation requires sufficient energy translocation across the inner plastid envelope

Low temperatures result in photoinhibition and consequently cold acclimation is accompanied by a limited plastidic ATP synthesis (Khanal et al., 2017). However, cold induced adaptations of thylakoid proteins, pigments or inner envelope composition essentially rely on sufficient ATP availability. NTT type carriers of higher plants act as ATP/ADP transporters and were shown to mediate energy provision to heterotrophic plastids as well as to autotrophic chloroplasts under conditions of missing or reduced photosynthetic activity (Reinhold et al., 2007; Kirchberger et al., 2008; Tjaden et al., 1998; Reiser et al., 2004). The comparative proteome study revealed that the abundance of NTT2 substantially increases in response to cold temperatures (log2FC +9.7, Table 1). Therefore, cold-induced limitations in photosynthetic energy production are apparently compensated by increased NTT-mediated ATP uptake from the cytosol. To test whether NTT activity is indeed required for proper cold acclimation we made use of NTT loss-of-function mutants. The Arabidopsis genome encodes two *ntt* isoforms and thus we analysed cold acclimation and acquisition of freezing tolerance in the corresponding single (*ntt1* and *ntt2*) mutants as well as in the double (*ntt1/2*) mutant (Reiser et al., 2004). After six weeks of growth at ambient conditions, *ntt1* and *ntt2* do not exhibit altered phenotypic appearance when compared to correspondingly grown wild type plants (WT). The double *ntt1/2* mutants however were slightly smaller (Figure 3B upper row).

**Figure 3:**
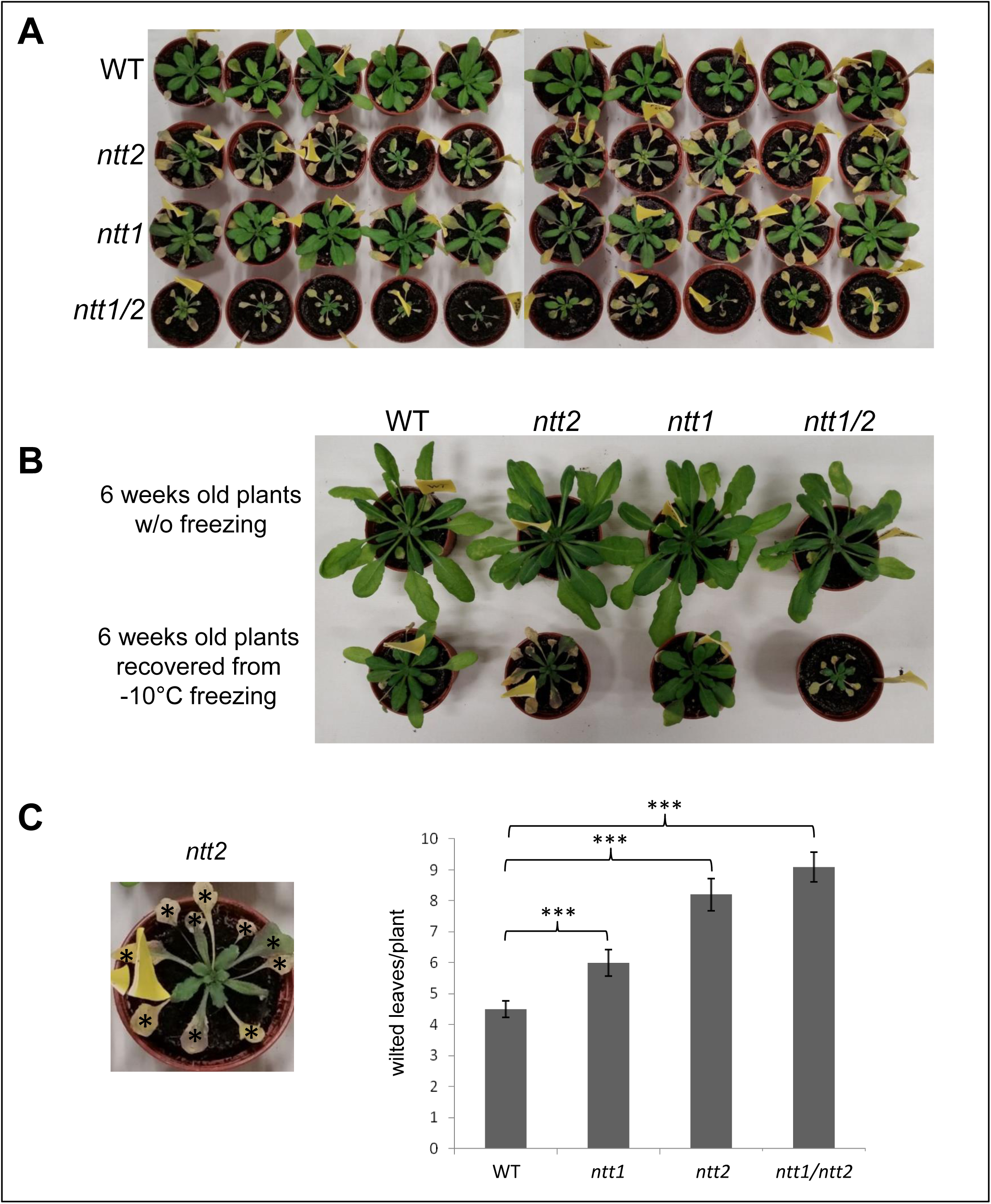
Effect of freezing to −10°C on WT (Col-0), *ntt2* T-DNA insertion mutant, *ntt1* T-DNA insertion mutant and the double *ntt1* and *ntt2* (ntt1/2) T-DNA insertion mutant. Plants were cultivated for 17 days under standard conditions (22°C day temperature, 18°C night temperature, day length 10h, relative humidity 60% and 120µE light intensity). Subsequent the temperature was lowered to 4°C day and night temperature (4 day cold acclimation) and afterwards the temperature was further lowered to −10°C (stepwise 2°C/hour). −10°C was kept for 15 hours before the temperature was raised again to 22°C (stepwise 2°C/hour). **A:** picture of WT, *ntt2*, *ntt1* and *ntt1/2* mutant plants recovered from −10°C freezing for 3 weeks. **B:** comparison of 6 week old WT, *ntt2*, *ntt1* and *ntt1/2* mutant plants with and without −10°C freezing treatment. **C:** quantification of wilted leaves from −10°C treated plants after 3 weeks recovery under standard growing conditions. The picture in C demonstrates in more detail how leaves were categorized as “wilted”. Statistics: n = 10, *** indicates a P value < 0.001 estimated by student’s t-test.

Moreover, the cold acclimation study revealed that after recovery from freezing, all three mutant lines exhibited more wilted leaves than the WT (Figure. 3A and C). The WT lost in average 4.2 leaves per plant, whereas *ntt1*, *ntt2* and *ntt1/*2 lost 6, 8.0 and 8.5 leaves per plant, respectively. The increased leave damage of plants lacking either NTT1 or NTT2 indicates that the activity of only one NTT isoform does not suffice to obtain proper freezing tolerance. Moreover, the observation that *ntt2* mutants exhibit more wilted leaves per plant than *ntt1* mutants and almost reach the number of dead leaves per plant of the double *ntt1/2* mutant suggests that NTT2 is of higher importance for cold acclimation than NTT1.

### Prevention of plastidic maltose export is required for proper freezing tolerance

It is well known that a tightly balanced cellular sugar- and starch homeostasis is critical for the plant’s capability to tolerate low or freezing temperatures (Nägele and Heyer, 2013; Pommerrenig et al., 2018). MEX1, the sole maltose exporter of the chloroplast was shown to play an important role in starch turnover and thus in the connection of starch and sugar metabolism (Purdy et al., 2013; Ryoo et al., 2013; Niittylä et al., 2004). Interestingly, cold exposure led to considerable depletion of this transport protein from the envelope proteome (log2FC −9.0, Table 1). Moreover, leaves of MEX1 loss-of-function mutants (*mex1-1*) were shown to exhibit metabolic features of cold acclimation already in the warm (Purdy et al., 2013). These observations imply that elevated maltose levels in the plastid are required for proper cold acclimation. Consequently, a constantly high maltose export activity might cause perturbations in cold acclimation. To test this hypothesis, we generated mutant plants overexpressing *mex1* and analysed their capacity to cope with freezing temperatures. For this, *mex1-1* mutants (Niittylä et al., 2004) were transformed with an expression construct carrying the structural *mex1* gene under control of the ubiquitin 10 promotor. Two strong overexpressor lines, *pUBQ10::MEX1* lines 1 and 2 (termed *pUBQ10::MEX1-1* and *pUBQ10::MEX1-2* respectively) were chosen for further studies (Supp. Fig. S2)

As previously shown, *mex1-1* mutants are highly impaired in growth when compared to the WT (Supp. Fig. S2) (Purdy et al., 2013; Niittylä et al., 2004) The two *mex1* overexpressor lines however grew much larger than *mex1-1* and showed WT appearance (Supp. Fig. S2). The fact that overexpressing mex1 complemented the dwarf phenotype of the original *mex1-1* mutant demonstrates that the introduced maltose transporter is functional (Supp. Fig. S2).

The cold acclimation study revealed that, although massively impaired in growth, the *mex1-1* mutant recovers quite well from freezing (Figure 4A). For quantitative evaluation of the freezing damage, we counted the wilted leaves of the individual plants. While WT plants exhibit 2.9 wilted leaves per plant in the mean, the number is only marginally increased (3.5 in the mean) in *mex1-1* plants (Figure 4B). By contrast, the two mex1 overexpressor lines are much more affected (Figure 4B) and show a significantly higher amount of wilted leaves per plant than the WT or *mex1-1*. An average of 6.3 and 5.8 leaves per plant of the *pUBQ10::MEX1-1* line and *pUBQ10::MEX1-2* line wilted from freezing, respectively (Figure 4). This result demonstrates that the overexpression of *mex1* results in higher susceptibility of the plants to cold stress and supports the idea that elevated maltose levels inside the chloroplast are required for cold acclimation.

**Figure 4:**
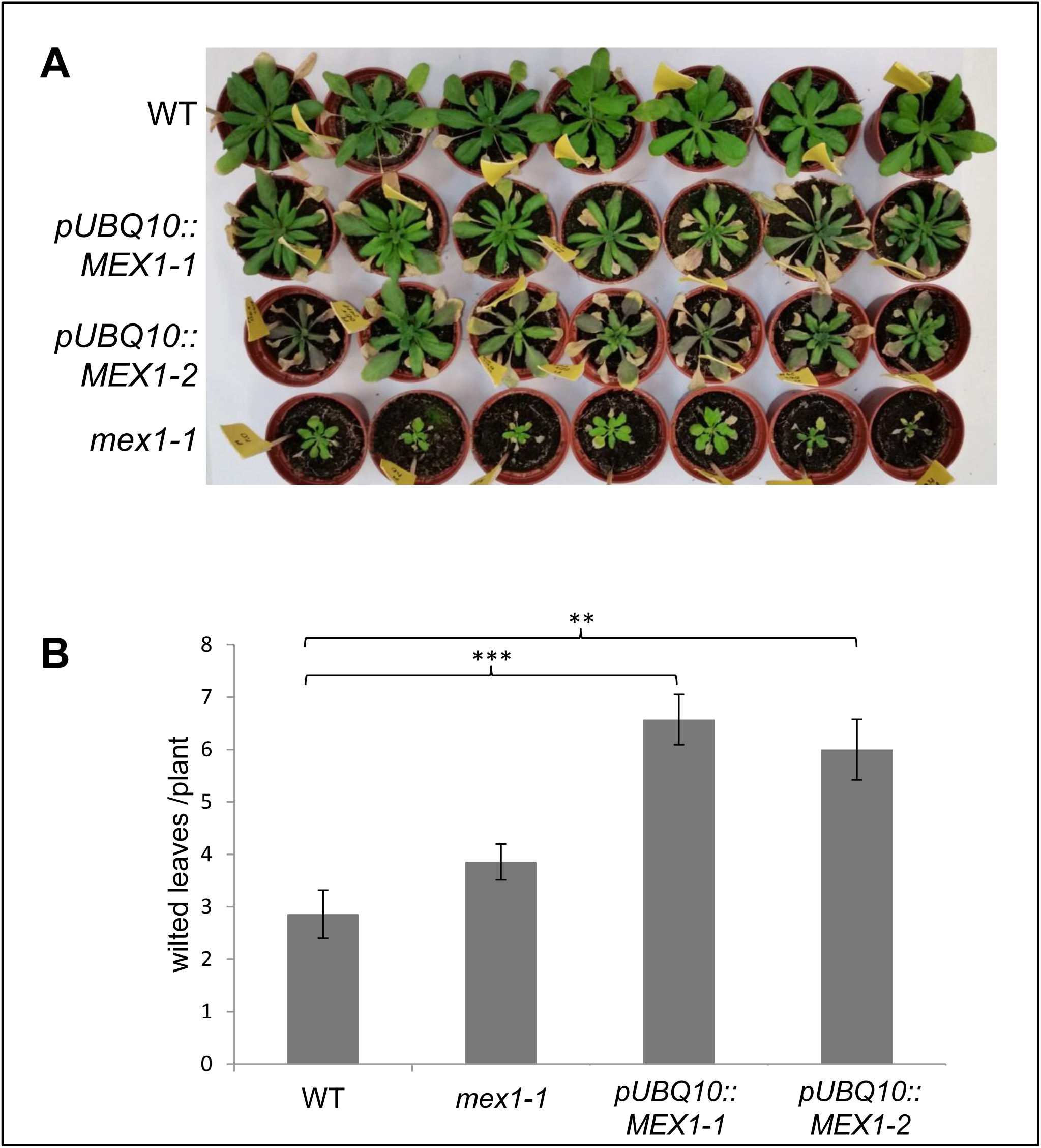
Effect of freezing to −10°C on WT (Col-0), *mex1-1* loss of function mutation and the *mex* overexpressor plants *pUBQ10::MEX1-1* and *pUBQ10::MEX1-2.* Plants were cultivated for 17 days under standard conditions (22°C day temperature, 18°C night temperature, day length 10h, relative humidity 60% and 120µE light intensity). Subsequent the temperature was lowered to 4°C day and night temperature (4 day cold acclimation) and afterwards the temperature was further lowered to −10°C (stepwise 2°C/hour). −10°C was kept for 15 hours before the temperature was raised again to 22°C (stepwise 2°C/hour). A: picture of WT, *mex1-1* mutation and overexpressor plants recovered from −10°C freezing for 3 weeks. B: quantification of wilted leaves from −10°C treated plants after 3 weeks recovery under standard growing conditions. Statistics: n = 7, *** indicates a P value < 0.001 and ** a P value < 0.01 estimated by student’s t-test.

### Assessing the cold acclimation dependent differentially localization of envelope associated soluble proteins

By using ratio-metric measurements comparing protein-specific enrichment factors under normal and cold conditions of the envelope (sub)-proteome (Supplement Table S1), we were able to access cold dependent differentially localization by mass spectrometry. Proteins with a positive log2FC change determined by the factor of their enrichment exhibit a higher abundance in the envelope fraction under cold treatment and vice versa. Superimposing the information about envelope localization for proteins that do not have any known transmembrane helices or other established transmembrane domains, dynamic differential localization due to cold becomes apparent (Figure 5 and Table 2). Therefore, an abundance increase determined by enrichment factor under cold conditions points to a conditional association of the proteins to the envelope membrane while a decrease in abundance suggests membrane dissociation. In total we were able to identify 24 non-intrinsic envelope proteins exhibiting a cold-acclimation dependent differential localization (q-value ≤ 0.05) without changes in abundance (Table 2). For three of these the changes are rather small and may have no biological relevance (Figure 5 grey dots) even though these changes are statistically relevant.

**Figure 5:**
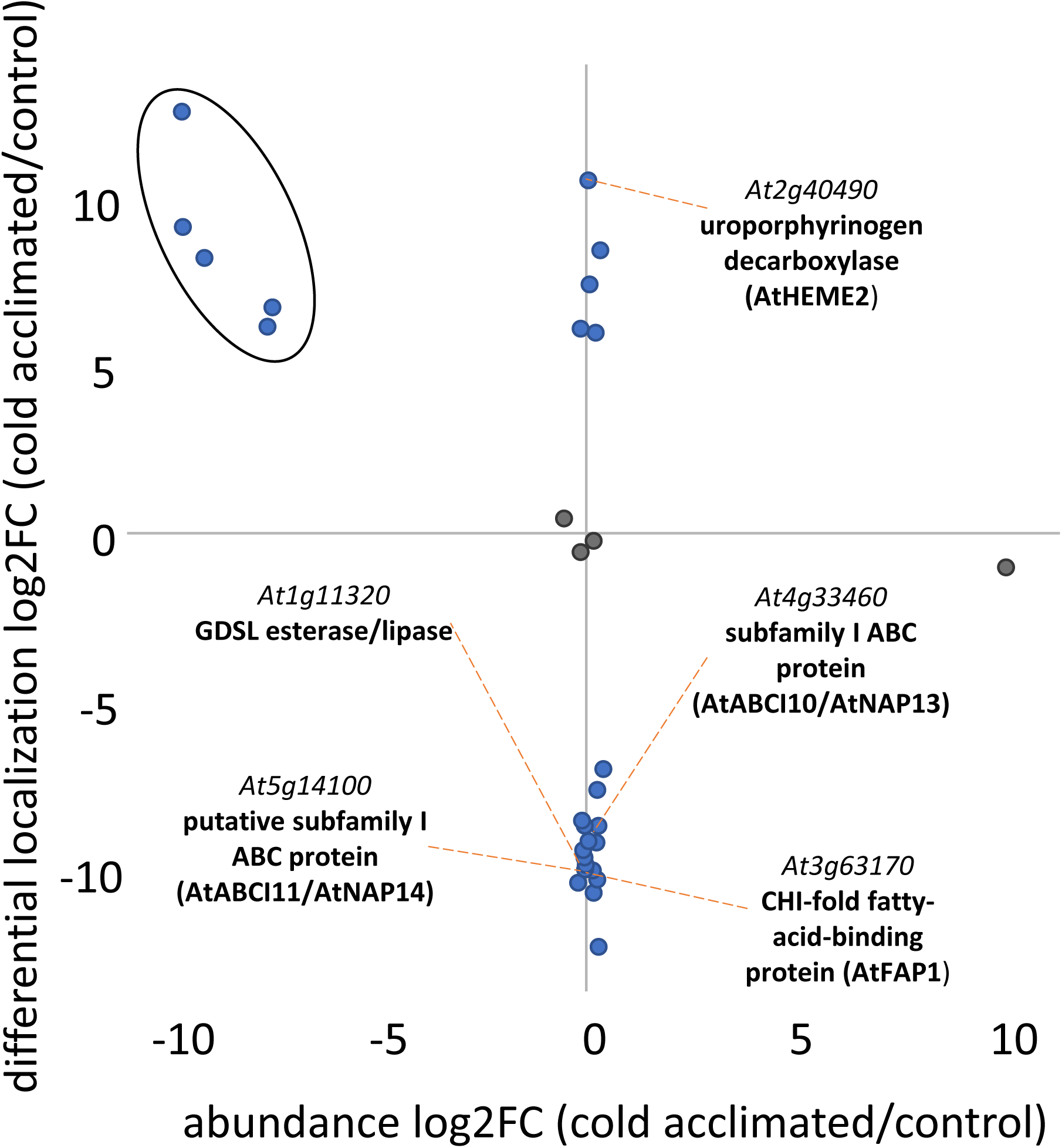
Cold dependent differentially localization of non-intrinsic envelope membrane associated proteins accessed by mass spectrometry. Proteins with a positive log2FC change determined by the factor of their enrichment exhibit a higher abundance in the envelope fraction under cold treatment and vice versa. Proteins exhibiting a rather small differential localization are indicated by grey dots.

**Table 2:** Identified proteins exhibiting a differential localization (diffloc) at envelope membrane

| Gene ID | Protein names | diffloc | q value<br>diffloc |
| --- | --- | --- | --- |
| *At3g44380* | late embryogenesis abundant (LEA) hydroxyproline-rich glycoprotein family | -12,37 | 6,88E-10 |
| *At4g23430* | translocon at the inner envelope membrane of chloroplasts 32-IVa | -10.75 | 4,66E-09 |
| *At1g11320* | GDSL esterase/lipase | -10.35 | 2,54E-09 |
| *At5g51070* | chaperone component of Clp-type protease complex (ClpD/ERD1) | -10.25 | 2,57E-08 |
| *At5g55510* | putative chloroplast envelope translocase component (PRAT1.2/HP22) | -10.04 | 9,46E-09 |
| *At5g14100* | subfamily I ABC protein (ABCI11/NAP14)) | -9,94 | 1,21E-09 |
| *At3g63170* | putative CHI-fold fatty-acid-binding protein (FAP1) | -9.77 | 0,000000239 |
| *At4g33460* | putative subfamily I ABC protein (ABCI10/NAP13) | -9.59 | 1,14E-08 |
| *At1g66670* | proteolytic component of Clp-type protease core complex (ClpP3/nClpP3) | -9.37 | 8,48E-09 |
| *At3g23700* | putative S1-type protein of small ribosomal subunit | -9.23 | 6,78E-10 |
| *At5g45170* | haloacid dehalogenase-like hydrolase (HAD) superfamily protein | -9.19 | 0,000000163 |
| *At1g52670* | putative regulator of acetyl-CoA carboxylase complex (BADC2) | -8.68 | 4,77E-09 |
| *At3g19720* | chloroplast fission mediator (DRP5B/AtARC5) | -8.64 | 2,31E-08 |
| *At1g63610* | protein of unknown function | -8.58 | 0,000000104 |
| *At3g13470* | putative component of plastidial Cpn60 chaperonin complex (CPN60B2) | -7.67 | 1,27E-08 |
| *At1g76180* | Early Response to Dehydration (ERD14) | -7.06 | 0,000000172 |
| *At5g14740* | beta carbonic anhydrase (BCA2/AtCA2) | -0.56 | 0,026906612 |
| *At4g13010* | inner envelope quinone-oxidoreductase, lacking cleavable N-terminal transit peptide (ceQORH) | -0.25 | 0,049506466 |
| *At1g12410* | non-proteolytic component of Clp-type protease core complex (ClpR2/nClpP2) | 0.41 | 0,0016654 |
| *At3g10230* | lycopene beta-cyclase (LCY-B) | 5.99 | 0,000000312 |
| *At3g26085* | CAAX amino terminal protease family protein | 6.08 | 0,00000979 |
| *At5g64816* | protein of unknown function | 7.38 | 0,000000128 |
| *At4g34120* | protein of unknown function, contains CBS-type domain (LEJ1/CDCP1) | 8.43 | 7,71E-08 |
| *At2g40490* | uroporphyrinogen decarboxylase (HEME2) | 10.52 | 1,95E-09 |

Interestingly, beside these two groups, there is a cluster of 5 proteins (decreased abundance) with a strong correlation between the differential localization and the overall change in protein abundance under cold (Figure 5 cluster indicated by the oval circle and Table 3). It seems that those proteins compensate for the global protein reduction during cold treatment by associating to membrane following their biochemical binding equilibrium. However, proteins showing differential localization that does not occur with a change in abundance might be modulated in their functional capacity. Additionally, there is one single protein with a high increase in abundance and a compared to the other 5 proteins exhibits a rather small differential localization (diffloc – 1.07, Figure 5 grey dot).

**Table 3:** Identified proteins exhibiting a differential localization at the envelope but also a difference in total abundance

| Gene ID | Protein names | diffloc | q value<br>diffloc | log2-FC<br>4°C - 22°C |
| --- | --- | --- | --- | --- |
| AT3G20330 | aspartate carbamoyltransferase (ATCase) | 12,57 | 0,0009683 | -9,80 |
| AT3G10840 | putative alpha/beta-fold-type hydrolas | 9,056 | 0,00000719 | -9,742 |
| AT1G65410 | putative component of ER-to-thylakoid lipid transfer complex<br>(TGD3/ABCI13/NAP11) | 8,17 | 0,000191721 | -9,25 |
| AT3G32930 | protein of unknown function | 6,17 | 4,74E-08 | -7,72 |
| AT3G08740 | Elongation factor P (EF-P) family protein | 6,67 | 0,000000138 | -7,60 |
| AT5G64840 | putative subfamily F ABC protein (ABCF5/GCN5) | -1,07 | 0,009831283 | 10,07 |

Among the identified proteins exhibiting a cold induced differential localization at the envelope membrane are several interesting candidates (Figure 5). At least two of these can be assigned to fatty acid, lipid-metabolism/modification: the chalcone isomerase-(CHI)-fold fatty-acid binding protein FAP1 (*At3g63170*) and the GDSL esterase/lipase (*At1g11320*). In Arabidopsis out of five so far characterized CHI-fold proteins three of these locate to the chloroplast and have been identified as fatty acid binding proteins (FAPs). They are highly expressed during increased fatty acid storage and knockout plants show elevated α-linolenic acid levels (Ngaki et al., 2012). GSDL lipolytic enzymes belong to a family of lipid hydrolysis enzymes widely existing in bacteria and plants (Lai et al., 2017). According to our analysis both proteins seem to dissociate from the envelope membrane during cold acclimation. An analogous behaviour has been observed by us for two proteins belonging to the subfamily I of the ABC protein family (AtABCI11/AtNAP14 *At5g14100* and AtABCI10/AtNAP13 *At4g33460*). Both are so called soluble non-intrinsic ABC proteins consisting solely out of one nucleotide binding site (Sánchez-Fernández et al., 2001) and one of these, NAP14, has already been characterized as involved in transition metal homeostasis. Disruption of the *nap14* gene results in over-accumulation of transition metals (Fe, Co, Cu, Zn and Mo) and abnormal chloroplast structures (Shimoni-Shor et al., 2010). However, the physiological role of NAP13 is so far unknown.

One of the proteins that exhibiting an increased attachment to the envelope membrane after cold is part of the tetrapyrrole pathway generating chlorophyll (*At*HEME2 *At2g40490*). The enzyme uroporphyrinogen decarboxylase (UROD) converts uroporphyrinogen III into coproporphyrinogen III which is than further converted to protophorphyrin IX (Terry and Smith, 2013). Depletion of UROD leads to reduction of the tetrapyrrole biosynthesis and light dependent necrosis (Mock and Grimm, 1997).

## Discussion

Our proteome study revealed cold-induced changes in the abundance of 38 envelope proteins. Although only three of these proteins increased in abundance, the corresponding alterations were considerably high and exhibited log2FC of 7.4 to 10.1. Moreover, among the 35 proteins of decreased abundance, 11 were highly reduced whereas the remaining 24 showed only moderate changes. The extreme positive or negative FC values apparently result from the very low abundance of the corresponding envelope proteins either under control or cold conditions. Therefore, the detection limit of the mass spectrometer apparently causes a certain degree of overestimation of their abundance changes. Independent of a possible overestimation, the observation that the amount of a very low abundant protein substantially increased is indicative for its stimulated synthesis whereas the almost complete disappearance of a highly abundant protein implies its effective degradation during cold exposure. One might envision that the moderate reduction of 24 envelope proteins results from a general reduction in protein biosynthesis due to the cold temperatures. However, because most envelope proteins remained unaffected by the cold treatment, it is likely that the observed abundance changes, either moderate or massive, are specific. In the following, we discuss the role of several altered proteins in cold acclimation.

### Energy provision to the chloroplast is required for proper freezing tolerance

Cold exposure initially results in decreased photosynthesis, which limits energy production in the chloroplast (Khanal et al., 2017). Therefore, metabolic processes in the stroma rely on additional energy provision from the cytosol. The antiporters NTT1 and NTT2 generally mediate ATP uptake into the stroma and play a central role particularly when photosynthetic energy production is insufficient (short day conditions) or missing (heterotrophic plastids) (Reinhold et al., 2007; Tjaden et al., 1998). Therefore, the considerable cold-induced increase in the abundance of NTT2 (+ 9.7 of 2-fold, Table 1) might lead to a stimulated ATP uptake from the cytosol. Here we demonstrate that absence of NTT2 or of both NTTs (Reiser et al., 2004) results in a substantially higher sensitivity of the corresponding mutant plants to freezing (Figure 3). This observation demonstrates that, cold acclimation relies on NTT mediated ATP supply to the chloroplast. The increased abundance of NTT2 most likely guarantees the energization of metabolic processes required for cold acclimation, such as the generation of starch or fatty acid synthesis.

### Cold acclimation involves blockage of plastidic maltose export

Sugars are important cryoprotectants, allow vitrification of membranes after water removal and quench ROS efficiently, and thus fulfil important functions during cold acclimation and acquisition of freezing tolerance. It is well known that photosynthetic carbohydrate fixation is essential for sugar accumulation in response to cold (Wanner and Junttila, 1999). However, several reports suggest that also starch breakdown contributes to the development of freezing tolerance, particularly during the early phase of low temperature exposure (Kaplan et al., 2006; Yano et al., 2005). Starch degradation generally results in the release of maltooligosaccharides and finally maltose. Although starch degradation seems to be prerequisite, it remained unclear whether the released maltose or maltose-derived compounds mediate proper cold acclimation and cryoprotection of the photosynthetic electron transport chain (Kaplan et al., 2006; Yano et al., 2005).

Maltose generally leaves the chloroplast via the transporter MEX1 (Niittylä et al., 2004). Interestingly, cold exposure led to considerable depletion of MEX1 from the envelope membrane (−9.0 log2FC, Table 1). This observation points to a prevention of maltose export from the plastid during cold temperatures. To gain deeper insights into the role of plastidic or cytosolic maltose in cold acclimation we generated plants constantly overexpressing MEX1. The reduced tolerance of the corresponding mutants against freezing (Figure 4) suggests that not only maltose release from starch degradation but also its trapping in the chloroplast is required for acquisition of proper freezing tolerance. This conclusion is supported by the fact that, although cellular maltose levels rise in response to high and low temperatures solely at low temperatures maltose accumulates in the chloroplast (Kaplan and Guy, 2004; Lu and Sharkey, 2006).

### Cold-induced changes in the lipid transfer across the chloroplast envelope

Cold exposure led to substantial changes in the relative abundance of several proteins associated with fatty acid (FA) and lipid metabolism. In fact, the modification of the chloroplast lipid composition is a conditio sine qua non for the acclimation and adaptation to low temperatures (Barrero-Sicilia et al., 2017; Li-Beisson et al., 2010). The synthesis of glycerolipids, which represent the major lipid constituents of plant membranes, takes place in two different compartments (Li-Beisson et al., 2010). The prokaryotic type of glycerolipid synthesis in the chloroplast gives rise to lipids almost exclusively carrying C16 fatty acids at the sn-2 position of the glycerol backbone, whereas the eukaryotic pathway at the endoplasmic reticulum generally produces glycerolipids with C18 fatty acids at the sn-2 position (Li-Beisson et al., 2010). In plants, the *de novo* synthesis of fatty acids however is restricted to the chloroplast and thus fatty acids must leave the organelle for modification at the ER. Moreover, the resulting ER-derived glycerolipids must enter the chloroplast. FAX1 catalyses fatty acid export from the chloroplast and consequently, mutant lines lacking this transporter exhibit decreased levels of ER-derived and increased levels of plastid-derived lipids (Li et al., 2015a). By contrast, the trigalactosyldiacylglycerol (TGD) protein complex mediates ER-to-chloroplast lipid transport (Roston et al., 2012). Moreover, missing or reduced presence of the substrate recognition domain TGD2 or the nucleotide binding domain TGD3 were shown to hamper translocation across the TGD complex (Lu and Benning, 2009; Lu et al., 2007). Thus, the cold-induced decrease of FAX1 might retain fatty acids in the chloroplast and by this stimulate the prokaryotic pathway whereas the cold-induced decrease of TGD2 and TGD3 decreases the uptake of lipids derived from the eukaryotic pathway (Table 1). In fact, cold temperatures were shown to be accompanied by increased carbon fluxes via the prokaryotic pathway and reduced contribution of the eukaryotic pathway (Li et al., 2015b). Therefore, we consider the lowering of the FAX1, TGD2 and TGD3 protein levels as a novel factor involved in cold-induced modulation of the glycerolipid composition of chloroplast membranes (Table 1).

### Cold acclimation is accompanied by changes in vesicle transfer at the chloroplast

Rab-GTPases fulfil different functions in vesicle transport and may be involved in vesicle budding, motility, tethering and docking. Interestingly, the abundance of the putative Rab-B-class GTPase RAB-B1b was shown to increase after cold treatment (Table 1). This protein is based on n-terminal YFP-(yellow fluorescent protein)-fusions predicted to localize to the secretory pathway (Chow et al 2008; Camacho et al. 2009) but was also found in the envelope membrane (Ferro et al., 2010; Bruley et al., 2012; Bouchnak et al. 2019) (Table 1). The cold-induced increase of RAB-B1b in the chloroplast fraction might thus be indicative for an enhanced fusion of ER-derived vesicles with the outer envelope. The corresponding vesicles might deliver new lipids or other cargos required for cold-induced changes of the chloroplast envelope. However, it is also imaginable that RAB-B1b is part of the intraplastidic vesicle trafficking system, which contributes to the modulation of the thylakoid membrane. Interestingly, cold treatment results in an increase in the lipid to protein ratio (Chapman et al., 1983) and is accompanied by an accumulation of vesicles in the stroma (Morré et al., 1991; Westphal et al., 2001). Moreover, already two RAB-GTPases (CPRabA5E, *At1g05810* and CPSAR1, *At5g18570*), have been identified to be involved in vesicle transport from the inner envelope to the thylakoid (Bang et al., 2009; Chigri et al., 2009; Garcia et al., 2010) and thus one might envision that RAB-B1b contributes to the cold-induced modulation of the thylakoid lipid content.

### Cold induced changes in the envelope point to alterations in nucleotide synthesis

In plants, the first steps of pyrimidine nucleotide *de novo* synthesis take place in the plastid stroma (Witz et al., 2012). The enzyme aspartate transcarbamylase (ACTase) (Hemsley et al., 2013) catalyses the second step in this biosynthesis pathway (Chen and Slocum, 2008). Interestingly, its lipid anchor and our proteome analysis suggest that the ATCase is attached to the inner envelope membrane, at least temporarily. The considerable decrease of the ACTase (*At3g20330*) abundance (Table 1) in the cold might limit *de nov*o synthesis of pyrimidine nucleotides. Moreover, the transporter (BT1; *At4g32400*), that delivers newly synthesised adenine nucleotides to the cytosol, shows decreased abundance in cold treated plants (Table 1). These observations imply that under cold conditions, the *de novo* synthesis of pyrimidine and purine nucleotides is of minor importance and that the corresponding salvage pathways might be enough to satisfy the cellular nucleotide demand. The lesser energy costs of these cytosolic salvage pathways (Witz et al. 2012) might represent an advantage particularly under cold conditions.

### Tocopherol synthesis is apparently altered in cold acclimated plants

The exposure of plants to low temperatures initially results in the overreduction of the electron transport chain and increased generation of ROS (Erling Tjus et al., 1998). To prevent the ROS-induced damage of fatty acids, plants synthesize specific antioxidants, the tocopherols (vitamin E) (Maeda et al., 2006; Munné-Bosch, 2002). At first glance the moderate decrease of two envelope enzymes of the vitamin E synthesis pathway, VTE6 and VTE3 (Table 1), appears to be contradictory to the importance of tocopherols in cold acclimation. However, in this context it is important to note that defects in tocopherol synthesis may cause changes in the composition of the individual vitamin E vitamers in Arabidopsis (Mène-Saffrané, 2017). Therefore, the moderate decrease in the abundance of VTE3 and VTE6 might represent a putative fine-tuning mechanism, shifting tocopherol biosynthesis towards the production of tocotrienols or PC-8, vitamers with even higher anti-oxidative function than α-tocopherol (Serbinova et al., 1991; Olejnik et al., 1997).

### Impact of cold acclimation on chlorophyll turnover

Various observations suggest that chlorophyll biosynthesis is strongly inhibited by cold temperatures (Tewari and Tripathy, 1998; Tewari and Tripathy, 1999). Interestingly, we identified that two envelope proteins involved in chlorophyll biosynthesis (Table 1) are of decreased abundance in the cold. The first one is the protoporphyrinogen IX oxidase 2 (PPO2 *At5g14220*), which catalyses the oxidation of protoporphyrinogen to protoporphyrin IX (Terry and Smith, 2013). The second one is a chaperone-like protein (CPP1 *At5g23040*) required for stabilization of the light-dependent protochlorophyllide oxidoreductase POR (Lee et al., 2013). Therefore, the cold-induced inhibition of chlorophyll biosynthesis becomes visible also on the level of the chloroplast envelope proteome. However, photosynthetic activity, an important prerequisite for cold acclimation, relies on the availability of chlorophyll. In this context, it is important to mention that TIC55, a part of the translocon of the inner membrane, shows decreased abundance in the cold (Table 1). Absence of TIC55 was recently shown to prevent chlorophyll degradation after induction of senescence (Chou et al., 2018). Therefore, the decreased abundance of TIC55 might help to keep a certain chlorophyll level during cold-induced inhibition of chlorophyll synthesis.

### Cold acclimation induces a differentially localization of envelope associated soluble proteins

A fundamental principle to regulate enzyme activities is to increase or decrease their cellular amounts. Beyond that, higher order regulations are post-translational modifications like e.g. protein phosphorylation, acetylation, n-linked glycosylation etc. However, when protein activities at or in a membrane must be modified a regulative dissociation or association of soluble proteins to/from the respective membrane might be imaginable.

Comparing the abundance of soluble proteins enriched with the purified envelopes membranes allowed us to identify proteins that are higher apparent or lesser apparent at the envelope membrane after 4 days of cold acclimation. Therefore, a negative value indicates a putative dissociation and a positive value an association from/to the envelope membrane (Figure 5, Table 2). We named this analysis differential localization (diffloc) and should not be mixed with a dual targeting (e.g. proteins targeted to mitochondria and chloroplast). Of course, such analyses only make sense for non-membrane intrinsic proteins (no TM domains or other membrane spanning domains like β-barrels, e.g. of OEPs). Additionally, putative candidates should not exhibit significant changes in total protein abundance at the level of total chloroplasts as this can result in negative or positive values not displaying a diffloc. This must be considered as diffloc values are calculated based on protein abundances.

We were able to exclude such candidates as the total abundance changes were determined based on the chloroplast samples. The vigorous treatment of the isolated envelope membranes with sodium carbonate diminishes the identification of weakly or better to say unspecific protein attachment to the envelope. Though, identification of envelope associated proteins without a biological relevance cannot completely be ruled out. Currently, it is only possible to speculate about the processes governing associating or dissociating of proteins and about their physiological relevance. However, it’s worth to mention that among the identified diffloc proteins there are several candidates exhibiting regulatory functions. For example, the three Clp proteins are members of the CLP protease system, a component of the chloroplast protease network (Olinares et al., 2011) essential for chloroplast development. Furthermore, the spectrum of processes in which the identified diffloc proteins are involved includes fatty acid metabolism (FAP1 and BADC1), lipid metabolism (GDSL esterase/lipase), carotenoid (LCY-B) and heme synthesis (HEME2), protein modification (CAAX amino terminal protease), protein translation (S1-type protein small ribosomal subunit) and several other.

In Figure 5 A cluster of proteins exhibiting high diffloc values is indicated which in addition exhibit high log2FC abundance decreases (compare Table 1). Despite this decrease in protein abundance latter proteins exhibit a positive diffloc value. This indicates that the association to the envelope membrane in the cold is a result of a marked biochemical equilibrium towards membrane binding. E.g., we assume that the identified association of the aspartate carbamoyltransferase (ATCase) reflects its prevalent localization at the chloroplast envelope.

## Conclusion

Our analyses revealed that the protein composition and content of the envelope membrane is apparently modified during cold acclimation. The abundance of most envelope membrane proteins was reduced and only three proteins increased in response to cold treatment. We selected two transport proteins and made use of corresponding mutant plants to analyse whether the observed abundance changes are relevant for cold acclimation. In fact, absence of the protein that usually increases during cold as well as a constantly high level of the protein that usually vanishes in the cold led to higher susceptibility of the plants to freezing. Furthermore, the identification of several proteins with known or postulated functions in fatty acid synthesis, lipid metabolism, and lipid protection is in line with the relevance of these membrane compounds in cold acclimation. The identified proteins are promising candidates for detailed future analyses unravelling their individual roles in cold acclimation and freezing tolerance. Our differential localisation analysis might display a new approach for the identification of transitional protein associations to the envelope.

## Materials and Methods

### Plant cultivation and cold acclimation conditions

*Arabidopsis thaliana* L. (ecotype Columbia, Col-0) were sown on standard soil (type ED73, Hermann Meyer KG-Germany, https://www.meyer-shop.com/) and stratified at 4°C for 24h in darkness. Afterwards, the plants were transferred to a plant cultivation chamber (Fitotron model SGR223, Weis Technik, Reiskirchen Germany). Cultivation condition: 22°C day temperature, 18°C night temperature, day length 10h, relative humidity 60% and 120µE light intensity. For cold acclimation, plants were incubated for 4 days at 4°C while all other cultivation parameters were kept constant. Non-acclimated plants were further cultivated under standard conditions as described above. Plant leaf material used for organelle purification was collected one hour before onset of light.

### Isolation of chloroplast envelope membranes

The envelope membrane isolation procedure can be divided into two steps: a) isolation of intact chloroplasts and b) enrichment of envelope membranes from these chloroplasts using a sucrose step gradient (Supp. Fig. S1). The isolation of intact chloroplasts was carried out with some modifications according an existing protocol (Kunst, 1998). 200 g leaf material were chopped off from 34 days old Arabidopsis plants (cold acclimated and control plants kept at 22°C) and transferred to ice-cold homogenization buffer medium (0.45 M sorbitol, 20 mM Tricine-KOH pH 8.4, 10 mM EDTA, 10 mM NaHCO_3_, 0.1% fatty-acid free bovine serum albumin). Ratio of buffer volume to weight of leaf material was 3:1 (v/w). In a glass-beaker the buffer/leaf mixture was further cooled in ice water to limit metabolic activity to a minimum. After 30 min the mixture was transferred to a 1 L stainless steel beaker. For a controlled rupture of the leaves, the blender was successively set on for 1s at setting “low”, 1 sec at “medium” and 1 sec at “high” (Waring blender commercial heavy-duty blender). This procedure was repeated twice. The disrupted leaf material was than filtered through 3 layers of Miracloth (http://www.merckmillipore.com), placed in a funnel and the flow through was collected in an ice cooled Erlenmeyer flask. From this suspension the chloroplast fraction was collected by centrifugation (1.000 *g*, 10 min, 4°C) and gently resuspended in 8 ml resuspension buffer medium (0.3M sorbitol, 20 mm Tricine-KOH, pH7.6, 5mM MgCl_2_, 2.5 mM EDTA), using a natural bristle paint brush.

A Percoll™ gradient was prepared by mixing equal volumes of 2-times concentrated resuspension buffer medium and pure Percoll^TM^. 30 ml of this mixture were transferred to 36 ml centrifuge tubes centrifuged (Sorval SS34 fixed angle rotor, 43.400 *g*, 30 min, 4°C, no brake). Two Percoll^TM^ gradients were enough for 200 g of leaf material. The Percoll gradient was overlaid with the resuspended chloroplast suspension. After centrifugation in a HB4 swing out rotor (13.300 *g*, 15 min, 4°C, no brake) two distinct green bands appeared (Supp. Fig.S1). The upper band, containing broken chloroplasts, was removed by using a water jet pump and the lower band was collected using a wide opened Pasteur pipette. This fraction was transferred to a SS34 tube and diluted with 3 volumes of resuspension buffer medium. From that suspension, intact chloroplasts were collected by centrifugation (HB4 rotor, 2.700 *g*, 5 min, no brake).

Enrichment of envelope membranes was carried with modifications according to a given protocol (Ferro et al., 2003). The intact chloroplast fraction was vigorously resuspended in 2 ml of buffer medium (10 mM MOPS-NaOH, pH 7.8) and kept on ice for 10 min to allow osmotic disruption of chloroplasts. To prevent protease driven protein degradation the buffer medium was complemented with a protease inhibitor cocktail (cOmplete™, EDTA-free, www.sigmaaldrich.comSigma). At this step 100 µl samples were collected from the lysate for the mass spectrometry-based identification of total chloroplast proteins.

A three-step sucrose gradient (bottom to top: 4 ml 0.93M, 0.6M and 0.3M sucrose) prepared in Ultra-Clear™ tubes (16×102 mm, www.beckmann.de) was overlaid with 1 ml of disrupted chloroplast preparation. After ultra-centrifugation (swing-out rotor SureSpin™ 630, www.thermo-fisher.com, 70.000 *g*, 1h, 4°C, no brake) the yellowish envelope fraction was collected from the inter-phase between 0.93M and 0.6M sucrose (Supp. Fig. S1). This fraction was 2-times diluted with double distilled water (_dd_H_2_O) and the envelope membranes were collected by ultra-centrifugation (ST120AT rotor, www.thermofisher.com, 400.000 *g*, 20 min, 4°C). The resulting membrane fraction was resuspended in 1ml _dd_H_2_O to remove any remaining sucrose. To remove membrane-associated proteins the envelope membranes were resuspended in 1M of sodium carbonate (Na_2_CO_3_) medium and centrifuged again, as described above. This washing step was repeated 5 times.

### Protein identification by tryptic digestion and mass spectrometry

Subsequently to protein estimation by Bradford assay (Bradford, 1976), the _dd_H_2_O resuspended envelope membranes or the total chloroplast samples membranes were solubilised by addition of SDS to a final volume of 2%, and 6-times concentrated SDS-Page loading dye was added (375mM Tris-HCl, pH6.8, 0.3% SDS (w/v), 60% glycerol (w/v), 1.5% bromphenolblue (w/v).

Equal amounts of envelope protein and chloroplast samples (isolated from cold acclimated or and control plants) were loaded on 12% SDS-Page. After Coomassie staining of the SDS-Page, the lanes were cut into 8 equal pieces and each piece was additionally cut into small cubes of approximately 1 mm side length. By consecutively (3-times) shrinking (with pure acetonitrile) and swelling (in 20 mM NH_4_HCO_3_) the buffer in which the gel pieces were resuspended, was removed before the proteins were reduced using 10mM DTT and alkylated using 55mM 2-iodoacetamide. Proteins were digested by addition of 12.5 ng/µl trypsin (Sigma-Pierce™, Trypsin Protease MS-Grade, www.thermoscientific.com) and incubation at 37°C for 15 h. Finally, peptides were extracted from the gel matrix using 2% trifluoroacetic acid. To guarantee a complete extraction of peptides the gel pieces were subsequent to a short centrifugation and removal of the supernatant shrunk again using acetonitrile. After short centrifugation the supernatants were collected, and the gel pieces were again treated with 2% trifluoroacetic acid. All three supernatants were pooled and dried down by vacuum centrifugation to approximately 30 µl. The peptide samples were desalted using handmade C18 STAGE tips following the protocol described by (Rappsilber et al., 2007). Finally, the C18 STAGE tip eluates were concentrated to approximately 2 µl and filled up to 20 µl with HPLC buffer A (2% acetonitrile, 0.1% formic acid).

### Protein identification and quantification

MS analysis was performed on a high-resolution LC-MS system (Eksigent nanoLC425 coupled to a Triple-TOF 6600, AB Sciex) in information dependent acquisition (IDA) mode. HPLC separation of 7.5 µl sample was performed in trap-elution mode using TriartC18 columns (5 µm particle, 0.5 × 5 mm for trapping and 3 µm particle, 300 µm × 150 mm for separation, YMC). A constant flow of 4 µl min^-1^ was employed and the gradient ramped within 15 min from 3 to 35% of HPLC buffer B (buffer A: 2% acetonitrile, 0.1% formic acid; buffer B: 90% acetonitrile, 0.1% formic acid), then within 1 min to 80% HPLC buffer B, followed by washing and equilibration steps. The mass spectrometer recorded one survey scan (250 ms accumulation time, 350–1250 m/z) and fragment spectra (100–1500 m/z) of the 30 most intense parent ions (30 ms accumulation time, charge state > 2, intensity > 300 cps, exclusion for 6 sec after one occurrence) resulting in a total cycle time of 1.2 sec. Identification and quantification of the proteins were performed using MaxQuant v. 1.6.0.16 (Cox and Mann, 2008). Spectra were matched against the ensemble plants release 43 of the Arabidopsis thaliana Tair10 genome release. The peptide database was constructed considering methionine oxidation and acetylation of protein N-termini as variable modifications and cabamido-methylation of cysteines as a fixed modification. False discovery rate (FDR) thresholds for peptide spectrum matches and protein identification were set to 1%. Protein quantification was carried out using the match-between-runs feature and the MaxQuant Label free Quantification (LFQ) algorithm (Cox et al., 2014). To make the complete mass spectrometric proteomic data available to the scientific community they have been deposited in the ProteomeXchange Consortium via the PRIDE partner repository (Perez-Riverol et al. 2018) with the dataset identifier PXD015794.

### Identification of cold regulated envelope proteins

In order to determine if a protein is localized to the chloroplast envelope and cold regulated, we decided to perform a multivariable logistic regression to integrate literature knowledge as well as data measured in this study. A positive and negative training data set was constructed using a combination of AT_CHLORO (Ferro et al., 2010; Bruley et al., 2012) and Plant Proteome Database (Kaplan et al., 2006; Sun et al., 2009) curated databases of sub-plastidial localization of proteins. As regressors we relied on a selected subset of protein sequence features defined in the AAindex1: Activation Gibbs energy of unfolding at pH 9.0, amino acid composition of MEM of single-spanning proteins, principal component II, hydrophobicity index, the Chou-Fasman parameter of coil conformation, average number of surrounding residues, interior composition of amino acids in intracellular proteins of mesophiles, weights for coil at the window position of −3, helix formation parameters, free energy in alpha-helical regions, average relative fractional occurrence in EL(i), and composition of amino acids in extracellular proteins (Zimmer et al., 2018). This set was extended using the experimental enrichment factors, calculated as the log2 fold change between the LFQ (abundances) of the plastid and envelope fractions. Subsequently, each score of the trained model was assigned to a posterior error probability (Käll et al., 2008). In the succeeding step to detect plastidic proteins with a differential abundance at low temperature treatment, we used the SAM method for statistical significance analysis (Larsson et al., 2005). Testing was performed using 4 biological replicates per condition. As response variables, the log2 transformed LFQ values of the plastidic fractions were used. A protein was treated as differentially regulated and envelope localized, if a q-value threshold of maximum 5% was not exceeded. Analyses was performed using Microsoft F# functional programming language with the bioinformatics library FSharpBio (available on GitHub: https://github.com/CSBiology/BioFSharp) in combination with the open source and cross-platform machine learning framework ML.NET. Charts were generated using the graphical chart library FSharp.Plotly (available on GitHub: https://github.com/muehlhaus/FSharp.Plotly). To assess the significance of differentially localizes proteins we compared the enrichment factors under cold and normal condition in the log2 space using Student’s T test statistic in SAM to account for multiple testing.

### Generation of AtMEX1 overexpression mutants

For cloning of *Atmex1*, gene sequences were amplified within a PCR reaction using the phusion-polymerase. For amplification a forward-primer containing a four base pair sequence (CACC) at its 5’end and a reverse-primer were used (*Atmex1*+1f_cacc: CACCATGGAAGGTAAAGCCATCGCG, AtMEX1c+1245r-stop: CGGTCCAAAAACAAGTTCTTTC). Cloning in the pENTR/D-Topo vector was done following the instruction of the pENTR™ Directional TOPO^®^ Cloning Kit (www.thermofisher.com/invitrogen). Entry vectors were then used to perform a recombination reaction with the expression-vector pUB-C-GFP (Grefen et al., 2010). The recombination reaction was done with the help of the GatewayTM LR ClonaseTM II Enzyme Mix (www.thermofisher.com/invitrogen) according to the guidelines of the manufacturer. *Atmex1* expression vectors were then used for heat-shock transformation of competent *Agrobacterium tumefaciens* cells (Höfgen and Willmitzer, 1988). Arabidopsis *mex1-1* mutant plants were then transformed according to a simplified version of the “floral dip method” suggested by Clough and Bent (Clough and Bent, 1998). Therefore, transformed Agrobacterium strains were grown in 200ml YEB liquid culture to an OD600 of approximately 0.8. Cells were harvested by centrifugation for 10 minutes with 4500 *g* at 4°C. 5% of sucrose (w/v) and 0.05% Silwet L77 (v/v) solved in water, were added to the Agrobacterium pellet. This mixture was then transferred in beakers. Five to six-week-old *Atmex1-1* plants, showing first closed inflorescences, were dipped in the bacteria culture for about 30 seconds. The dipped plants were transferred to plastic trays and were covered by a plastic hood for the following 48 hours. After 48 hours the plants were returned to their normal growing conditions and seeds were harvested three to four weeks after dipping. Seeds of Arabidopsis plants obtained from *Agrobacterium tumefaciens* mediated transformation were germinated on soil and selected by spraying with 0.1% BASTA (Glufosinat-Ammoniumsalt) herbicide (Logemann et al., 2006). Spraying was carried out on plants one week after germination and was repeated four times in intervals of two days.

### Freezing tolerance test

To perform a freezing tolerance test seeds of Arabidopsis WT and mutant plants were sowed and stratified as described above. After one-week seedlings were pricked into small pots containing standard soil (see above) and further cultivated in a Percival plant growth cabinet (Typ AR-36L/LT www.plantclimatics.de) under the described conditions. After further cultivation for 10 days the day and night temperature were lowered to 4°C for four days (cold acclimation). At the end of the night period of the fourth day the illumination was set of and the temperature was lowered from 4°C to −10°C in a stepwise manner (2°C per hour). The freezing temperature was kept constant for 15 hours before the temperature was increased to 22°C in a step wise manner (2°C per hour). Afterwards normal growing conditions were restored (see above). Plants were daily inspected and wilting of leaves was documented by photographs.

## Notes

Funding information: The work presented here was supported by the German Research Foundation (DFG) within the Collaborative Research Centre 175 (Transregio Sonderforschungsbereich 175)

## Literature cited

Ajjawi I, Coku A, Froehlich JE, Yang Y, Osteryoung KW, Benning C, Last RL (2011) A J-like protein influences fatty acid composition of chloroplast lipids in Arabidopsis. PLoS ONE 6 (10): e25368. doi: 10.1371/journal.pone.0025368

Alberdi M, Corcuera LJ (1991) Cold acclimation in plants. Phytochemistry (United Kingdom)

Amme S, Matros A, Schlesier B, Mock HP (2006) Proteome analysis of cold stress response in Arabidopsis thaliana using DIGE-technology. J Exp Bot 57: 1537–1546.

Awai K, Xu C, Tamot B, Benning C (2006) A phosphatidic acid-binding protein of the chloroplast inner envelope membrane involved in lipid trafficking. Proc Natl Acad Sci USA 103: 10817–10822

Bang WY, Hata A, Jeong IS, Umeda T, Masuda T, Chen J, Yoko I, Suwastika IN, Kim DW, Im CH, Lee BH, Lee Y, Lee KW, Shiina T, Bahk JD (2009) AtObgC, a plant ortholog of bacterial Obg, is a chloroplast-targeting GTPase essential for early embryogenesis. Plant Mol Biol 71: 379–390

Barrero-Sicilia C, Silvestre S, Haslam RP, Michaelson LV (2017) Lipid remodelling: Unravelling the response to cold stress in Arabidopsis and its extremophile relative *Eutrema salsugineum*. Plant Sci 263: 194–200

Bouchnak I, Brugière S, Moyet L, Le Gall S, Salvi D, Kuntz, Marcel K, Tardif M, Rolland N (2019) Unraveling Hidden Components of the Chloroplast Envelope Proteome: Opportunities and Limits of Better MS Sensitivity. MCP 18: 1285–1306.

Bradford MM (1976) A rapid and sensitive method for the quantitation of microgram quantities of protein utilizing the principle of protein-dye binding. Anal Biochem 72: 248–254

Bruley C, Dupierris V, Salvi D, Rolland N, Ferro M (2012) AT_CHLORO: A chloroplast protein database dedicated to sub-plastidial localization. Front Plant Sci 3: 205. doi: 10.3389/fpls.2012.00205

BB Buchanan (2015) Biochemistry & molecular biology of plants, Ed. 2. Wiley Blackwell; American Society of Plant Biologists, Chichester, Rockville, Md, USA

Calixto CPG, Guo W, James AB, Tzioutziou NA, Entizne JC, Panter PE, Knight H, Nimmo HG, Zhang R, Brown JWS (2018) Rapid and dynamic alternative splicing impacts the Arabidopsis cold response transcriptome. Plant Cell 30: 1424–1444

Camacho L, Smertenko AP, Pérez-Gómez J, Hussey PJ, Moore I (2009) Arabidopsis Rab-E GTPases exhibit a novel interaction with a plasma-membrane phosphatidylinositol-4-phosphate 5-kinase. J Cell Sci 122: 4383–4392

Catalá R, Medina J, Salinas J (2011) Integration of low temperature and light signalling during cold acclimation response in Arabidopsis. Proc Natl Acad Sci USA 108: 16475–16480

Chapman DJ, De-Felice J, Barber J (1983) Growth temperature effects on thylakoid membrane lipid and protein content of pea chloroplasts. Plant Physiol 72: 225–228

Chen CT, Slocum RD (2008) Expression and functional analysis of aspartate transcarbamoylase and role of de novo pyrimidine synthesis in regulation of growth and development in Arabidopsis. Plant Physiol Biochem 46: 150–159

Chen J, Han G, Shang C, Li J, Zhang H, Liu F, Wang J, Liu H, Zhang Y (2015): Proteomic analyses reveal differences in cold acclimation mechanisms in freezing-tolerant and freezing-sensitive cultivars of alfalfa. Front Plant Sci 6. doi: 10.3389/fpls.2015.00105.

Cheng Z, Sattler S, Maeda H, Sakuragi Y, Bryant DA, DellaPenna D (2003) Highly divergent methyltransferases catalyze a conserved reaction in tocopherol and plastoquinone synthesis in cyanobacteria and photosynthetic eukaryotes. Plant Cell 15: 2343–2356

Chigri F, Sippel C, Kolb M, Vothknecht UC (2009) Arabidopsis OBG-like GTPase (AtOBGL) is localized in chloroplasts and has an essential function in embryo development. Mol Plant 2: 1373–1383

Chou ML, Liao WY, Wei WC, Li AYS, Chu CY, Wu CL, Liu CL, Fu TH, Lin LF (2018) The direct involvement of dark-induced Tic55 protein in chlorophyll catabolism and its indirect role in the MYB108-NAC signalling pathway during leaf senescence in Arabidopsis thaliana. Int J Mol Sci 19: 1854; doi:10.3390/ijms19071854

Chow CM, Neto H, Foucart C Moore I (2008) Rab-A2 and Rab-A3 GTPases define a trans-golgi endosomal membrane domain in Arabidopsis that contributes substantially to the cell plate. Plant Cell 20: 101–123.

Clough SJ, Bent AF (1998) Floral dip: a simplified method for Agrobacterium-mediated transformation of Arabidopsis thaliana. Plant J 16: 735–743

Cox J, Hein MY, Luber CA, Paron I, Nagaraj N, Mann M (2014) Accurate proteome-wide label-free quantification by delayed normalization and maximal peptide ratio extraction, termed MaxLFQ. Mol Cell Proteom 13: 2513–2526

Cox J, Mann M (2008) MaxQuant enables high peptide identification rates, individualized p.p.b.-range mass accuracies and proteome-wide protein quantification. Nature Biotechnol 26: 1367–1372

Crosatti C, Rizza F, Badeck FW, Mazzucotelli E, Cattivelli L (2013) Harden the chloroplast to protect the plant. Physiol Plant 147: 55–63

Erling Tjus S, Lindberg Møller B, Vibe Scheller H (1998) Photosystem I is an early target of photoinhibition in barley illuminated at chilling temperatures. Plant Physiol 116: 755–764.

Ferro M, Brugière S, Salvi D, Seigneurin-Berny D, Court M, Moyet L, Ramus C, Miras S, Mellal M, Le Gall S, Kieffer-Jaquinod S, Bruley C, Garin J, Joyard J, Masselon C, Rolland N (2010) AT_CHLORO, a comprehensive chloroplast proteome database with subplastidial localization and curated information on envelope proteins. Mol Cell Proteom 9: 1063–1084

Ferro M, Salvi D, Brugière S, Miras S, Kowalski S, Louwagie M, Garin J, Joyard J, Rolland N (2003) Proteomics of the chloroplast envelope membranes from Arabidopsis thaliana. Mol Cell Proteom 2: 325–345

Fritsche S, Wang X, Jung C (2017) Recent advances in our understanding of tocopherol biosynthesis in plants: An overview of key genes, functions, and breeding of Vitamin E improved crops. Antioxidants (Basel) 6: E99. doi: 10.3390/antiox6040099

Gao F, Zhou Y, Zhu W, Li X, Fan L, Zhang G (2009): Proteomic analysis of cold stress-responsive proteins in Thellungiella rosette leaves. Planta 230: 1033–1046

Garcia C, Khan NZ, Nannmark U, Aronsson H (2010) The chloroplast protein CPSAR1, dually localized in the stroma and the inner envelope membrane, is involved in thylakoid biogenesis. Plant J 63: 73–85

Goetze TA, Patil M, Jeshen I, Bölter B, Grahl S, Soll J (2015) Oep23 forms an ion channel in the chloroplast outer envelope. BMC Plant Biol 15: 47. doi: 10.1186/s12870-015-0445-1

Goulas E, Schubert M, Kieselbach T, Kleczkowski LA, Gardeström P, Schröder W, Hurry V (2006): The chloroplast lumen and stromal proteomes of Arabidopsis thaliana show differential sensitivity to short- and long-term exposure to low temperature. Plant J 47: 720–734.

Grefen C, Donald N, Hashimoto K, Kudla J, Schumacher K, Blatt MR (2010) A ubiquitin-10 promoter-based vector set for fluorescent protein tagging facilitates temporal stability and native protein distribution in transient and stable expression studies. Plant J 64: 355–365

Haferkamp I, Schmitz-Esser S (2012) The plant mitochondrial carrier family: functional and evolutionary aspects. Front Plant Sci 3: 2 doi: 10.3389/fpls.2012.00002

Hemsley PA, Weimar T, Lilley KS, Dupree P, Grierson CS (2013) A proteomic approach identifies many novel palmitoylated proteins in Arabidopsis. New Phytol 197: 805–814

Hincha DK (2008) Effects of alpha-tocopherol (vitamin E) on the stability and lipid dynamics of model membranes mimicking the lipid composition of plant chloroplast membranes. FEBS Lett 582: 3687–3692

Höfgen R, Willmitzer L (1988) Storage of competent cells for Agrobacterium transformation. Nucleic Acids Res 16: 9877

Käll L, Storey JD, MacCoss MJ, Noble WS (2008) Posterior error probabilities and false discovery rates: two sides of the same coin. J Proteome Res 7: 40–44

Kampfenkel K, Möhlmann T, Batz O, van Montagu M, Inzé D, Neuhaus HE (1995) Molecular characterization of an Arabidopsis thaliana cDNA encoding a novel putative adenylate translocator of higher plants. FEBS lett 374: 351–355

Kaplan F, Guy CL (2004) β-Amylase induction and the protective role of maltose during temperature shock. Plant Physiol 135: 1674–1684

Kaplan F, Sung DY, Guy CL (2006) Roles of beta-amylase and starch breakdown during temperatures stress. Physiol Planta 126: 120–128

Karim S, Aronsson H (2014) The puzzle of chloroplast vesicle transport – involvement of GTPases. Front Plant Sci 5: 472 doi: 10.3389/fpls.2014.00472

Khanal N, Bray GE, Grisnich A, Moffatt BA, Gray GR (2017) Differential mechanisms of photosynthetic acclimation to light and low temperature in Arabidopsis and the extremophile Eutrema salsugineum. Plants (Basel) 6: (3). pii: E32. doi: 10.3390/plants6030032

Kim H, Botelho SC, Park K, Kim H (2015) Use of carbonate extraction in analyzing moderately hydrophobic transmembrane proteins in the mitochondrial inner membrane. Protein Sci 24: 2063–2069

Kirchberger S, Tjaden J, Neuhaus HE (2008) Characterization of the Arabidopsis Brittle1 transport protein and impact of reduced activity on plant metabolism. Plant J 56: 51–63

Kleine T, Leister D (2013) Retrograde signals galore. Front Plant Sci 4: 45 doi: 10.3389/fpls.2013.00045

Knaupp M, Mishra KB, Nedbal L, Heyer AG (2011) Evidence for a role of raffinose in stabilizing photosystem II during freeze-thaw cycles. Planta 234: 477–486

Kosová K, Vítámvás P, Planchon S, Renaut J, Vanková R, Prášil IT (2013) Proteome analysis of cold response in spring and winter wheat (*Triticum aestivum*) crowns reveals similarities in stress adaptation and differences in regulatory processes between the growth habits. J Proteome Res 12: 4830–4845.

Kötting O, Kossmann J, Zeeman SC, Lloyd JR (2010) Regulation of starch metabolism: the age of enlightenment? Curr Opin Plant Biol 13: 321–329

Kunst L (1998) Preparation of physiologically active chloroplasts from Arabidopsis. Methods Mol Biol 82: 43–48

Lai CP, Huang LM, Chen LFO, Chan MT, Shaw JF (2017) Genome-wide analysis of GDSL-type esterases/lipases in Arabidopsis. Plant Mol Biol 95: 181–197

Larsson O, Wahlestedt C, Timmons JA (2005) Considerations when using the significance analysis of microarrays (SAM) algorithm. BMC Bioinform 6: 129

Lee JY, Lee HS, Song JY, Jung YJ, Reinbothe S, Park YI, Lee SY, Pai HS (2013) Cell growth defect factor1/chaperone-like protein of POR1 plays a role in stabilization of light-dependent protochlorophyllide oxidoreductase in Nicotiana benthamiana and Arabidopsis. Plant Cell 25: 3944–3960

Li N, Gügel IL, Giavalisco P, Zeisler V, Schreiber L, Soll J, Philippar K (2015a) FAX1, a novel membrane protein mediating plastid fatty acid export. PLoS Biol 13: e1002053

Li Q, Zheng Q, Shen W, Cram D, Fowler DB, Wei Y, Zou J (2015b) Understanding the Biochemical Basis of Temperature-Induced Lipid Pathway Adjustments in Plants. Plant Cell 27: 86–103

Li-Beisson Y, Shorrosh B, Beisson F, Andersson MX, Arondel V, Bates PD, Baud S, Bird D, Debono A, Durrett TP, Franke RB, Graham IA, Katayama K, Kelly AA, Larson T, Markham JE, Miquel M, Molina I, Nishida I, Rowland O, Samuels L, Schmid KM, Wada H, Welti R, Xu C, Zallot R, Ohlrogge J (2010) Acyl-lipid metabolism. The Arabidopsis book 8: e0133

Logemann E, Birkenbihl RP, Ülker B, Somssich IE (2006) An improved method for preparing Agrobacterium cells that simplifies the Arabidopsis transformation protocol. Plant Methods 2: 16 doi: 10.1186/1746-4811-2-16

Lu B, Benning C (2009) A 25-Amino Acid Sequence of the Arabidopsis TGD2 Protein Is Sufficient for Specific Binding of Phosphatidic Acid. J Biol Chem 284: 17420–17427

Maeda H, Song W, Sage TL, DellaPenna D (2006) Tocopherols play a crucial role in low-temperature adaptation and Phloem loading in Arabidopsis. Plant Cell 18: 2710–2732

Maurer-Stroh S, Eisenhaber F (2005) Refinement and prediction of protein prenylation motifs. Genome Biol 6: R55

McFadden GI (1999) Endosymbiosis and evolution of the plant cell. Curr Opin Plant Biol 2: 513–519

Mène-Saffrané L (2017) Vitamin E biosynthesis and its regulation in plants. Antioxidants (Basel) 7(1) pii: E2. doi: 10.3390/antiox7010002

Mock HP, Grimm B (1997) Reduction of uroporphyrinogen decarboxylase by antisense RNA expression affects activities of other enzymes involved in tetrapyrrole biosynthesis and leads to light-dependent necrosis. Plant Physiol 113: 1101–1112

Moellering ER, Muthan B, Benning C (2010) Freezing tolerance in plants requires lipid remodeling at the outer chloroplast membrane. Science 330: 226–228

Morré DJ, Selldén G, Sundqvist C, Sandelius AS (1991) Stromal low temperature compartment derived from the inner membrane of the chloroplast envelope. Plant Physiol 97: 1558–1564

Munné-Bosch S (2002) The function of tocopherols and tocotrienols in plants. Crit Rev Plant Sci 21: 31–57

Nägele T, Heyer AG (2013) Approximating subcellular organisation of carbohydrate metabolism during cold acclimation in different natural accessions of *Arabidopsis thaliana*. New Phytol 198: 777–787

Ngaki MN, Louie GV, Philippe RN, Manning G, Pojer F, Bowman ME, Li L, Larsen E, Wurtele ES, Noel JP (2012) Evolution of the chalcone-isomerase fold from fatty-acid binding to stereospecific catalysis. Nature 485: 530–533

Niittylä T, Messerli G, Trevisan M, Chen J, Smith AM, Zeeman SC (2004) A previously unknown maltose transporter essential for starch degradation in leaves. Science 303: 87–89

Olejnik D, Gogolewski M, Nogala-Kałucka M (1997) Isolation and some properties of plastochromanol-8. Food 41: 101–104

Olinares PDB, Kim J, van Wijk KJ (2011) The Clp protease system; a central component of the chloroplast protease network. Biochim Biophys Acta 1807: 999–1011

Patzke K, Prananingrum P, Klemens PAW, Trentmann O, Rodrigues CM, Keller I, Fernie AR, Geigenberger P, Bölter B, Lehmann M, Schmitz-Esser S, Pommerrenig B, Haferkamp I, Neuhaus HE (2019) The plastidic sugar transporter pSuT influences flowering and affects cold responses. Plant Physiol 179: 569–587

Perez-Riverol Y, Csordas A, Bai J, Bernal-Llinares M, Hewapathirana S, Kundu DJ, Inuganti A, Griss J, Mayer G, Eisenacher M, Pérez E, Uszkoreit J, Pfeuffer J, Sachsenberg T, Yilmaz S, Tiwary S, Cox J, Audain E, Walzer M, Jarnuczak AF, Ternent T, Brazma A, Vizcaíno JA (2019) The PRIDE database and related tools and resources in 2019: improving support for quantification data. Nucleic Acids Res 47(D1): D442–D450

Pfalz J, Liebers M, Hirth M, Grübler B, Holtzegel U, Schröter Y, Dietzel L, Pfannschmidt T (2012) Environmental control of plant nuclear gene expression by chloroplast redox signals. Front Plant Sci 3:257. doi: 10.3389/fpls.2012.00257

Pommerrenig B, Ludewig F, Cvetkovic J, Trentmann O, Klemens PAW, Neuhaus HE (2018) In concert: orchestrated changes in carbohydrate homeostasis are critical for plant abiotic stress tolerance. Plant Cell Physiol 59: 1290–1299

Purdy SJ, Bussell JD, Nunn CP, Smith SM (2013) Leaves of the Arabidopsis maltose exporter1 mutant exhibit a metabolic profile with features of cold acclimation in the warm. PloS one 8 (11): e79412. doi: 10.1371/journal.pone.0079412

Rappsilber J, Mann M, Ishihama Y (2007) Protocol for micro-purification, enrichment, pre-fractionation and storage of peptides for proteomics using StageTips. Nature Prot 2: 1896–1906

Reinhold T, Alawady A, Grimm B, Beran KC, Jahns P, Conrath U, Bauer J, Reiser J, Melzer M, Jeblick W, Neuhaus HE (2007) Limitation of nocturnal import of ATP into Arabidopsis chloroplasts leads to photooxidative damage. Plant J 50: 293–304

Reiser J, Linka N, Lemke L, Jeblick W, Neuhaus HE (2004) Molecular physiological analysis of the two plastidic ATP/ADP transporters from Arabidopsis. Plant Physiol 136: 3524–3536

Rekarte-Cowie I, Ebshish OS, Mohamed KS, Pearce RS (2008) Sucrose helps regulate cold acclimation of Arabidopsis thaliana. J Exp Bot 59: 4205–4217

Ren J, Wen L, Gao X, Jin C, Xue Y, Yao X (2008) CSS-Palm 2.0: an updated software for palmitoylation sites prediction. Protein Eng Des Sel 21: 639–644

Rocco M, Arena S, Renzone G, Scippa GS, Lomaglio T, Verrillo F, Scaloni A, Marra M (2013): Proteomic analysis of temperature stress-responsive proteins in Arabidopsis thaliana rosette leaves. Mol Biosyst 9: 1257–1267

Röhl T, Motzkus M, Soll J (1999) The outer envelope protein OEP24 from pea chloroplasts can functionally replace the mitochondrial VDAC in yeast. FEBS lett 460: 491–494

Roston RL, Gao J, Murcha MW, Whelan J, Benning C (2012) TGD1, −2, and −3 proteins involved in lipid trafficking form ATP-binding cassette (ABC) transporter with multiple substrate-binding proteins. J Biol Chem 287: 21406–21415

Ryoo N, Eom JS, Kim HB, Vo BT, Lee SW, Hahn TR, Jeon JS (2013) Expression and functional analysis of rice plastidic maltose transporter, OsMEX1. J Korean Soc Appl Biol Chem 56: 149–155

Sánchez-Fernández R, Davies TG, Coleman JO, Rea PA (2001) The Arabidopsis thaliana ABC protein superfamily, a complete inventory. J Biol Chem 276: 30231–30244

Schneider T, Keller F (2009) Raffinose in chloroplasts is synthesized in the cytosol and transported across the chloroplast envelope. Plant Cell Physiol 50: 2174–2182

Schulze WX, Schneider T, Starck S, Martinoia E, Trentmann O (2012) Cold acclimation induces changes in Arabidopsis tonoplast protein abundance and activity and alters phosphorylation of tonoplast monosaccharide transporters. Plant J 69: 529–541

Serbinova E, Kagan V, Han D, Packer L (1991) Free radical recycling and intramembrane mobility in the antioxidant properties of alpha-tocopherol and alpha-tocotrienol. Free Radic. Biol. Med 10: 263–275

Sicher R (2011) Carbon partitioning and the impact of starch deficiency on the initial response of Arabidopsis to chilling temperatures. Plant Sci 181: 167–176.

Strand A, Hurry V, Gustafsson P, Gardeström P (1997) Development of Arabidopsis thaliana leaves at low temperatures releases the suppression of photosynthesis and photosynthetic gene expression despite the accumulation of soluble carbohydrates. Plant J 12: 605–614

Sun Q, Zybailov B, Majeran W, Friso G, Olinares PDB, van Wijk KJ (2009) PPDB, the plant proteomics database at cornell. Nucleic Acids Res 37: D969–74. doi: 10.1093/nar/gkn654.

Terry MJ, Smith AG (2013) A model for tetrapyrrole synthesis as the primary mechanism for plastid-to-nucleus signaling during chloroplast biogenesis. Front Plant Sci 4: 14. doi: 10.3389/fpls.2013.00014

Tewari KA, Tripathy CB (1998) Temperature-stress-induced impairment of chlorophyll biosynthetic reactions in cucumber and wheat. Plant Physiol 117: 851–858

Tewari KA, Tripathy CB (1999) Acclimation of chlorophyll biosynthetic reactions to temperature stress in cucumber (Cucumis sativus L.). Planta 208: 431–437

Tjaden J, Möhlmann T, Kampfenkel K, Neuhaus, Henrichs G and H. Ekkehard (1998) Altered plastidic ATP/ADP-transporter activity influences potato (Solanum tuberosum L.) tuber morphology, yield and composition of tuber starch. Plant J 16: 531–540

Trentmann O, Haferkamp I (2013) Current progress in tonoplast proteomics reveals insights into the function of the large central vacuole. Front Plant Sci 4: 34. doi: 10.3389/fpls.2013.00034

Trentmann O, Jung B, Neuhaus HE, Haferkamp I (2008) Nonmitochondrial ATP/ADP transporters accept phosphate as third substrate. J Bio Chem 283: 36486–36493

van Wijk KJ, Kessler F (2017) Plastoglobuli: Plastid microcompartments with integrated functions in metabolism, plastid developmental transitions, and environmental adaptation. Annu. Rev. Plant Bio 68: 253–289

Wanner LA, Junttila O (1999) Cold-induced freezing tolerance in Arabidopsis. Plant Physiol 120: 391–400

Weber A, Servaites JC, Geiger DR, Kofler H, Hille D, Gröner F, Hebbeker U, Flügge UI (2000) Identification, purification, and molecular cloning of a putative plastidic glucose translocator. Plant Cell 12: 787–802

Weber APM, Schwacke R, Flügge UI (2005) Solute transporters of the plastid envelope membrane. Annu. Rev. Plant Bio 56: 133–164

Westphal S, Soll J, Vothknecht UC (2001) A vesicle transport system inside chloroplasts. FEBS letters 506: 257–261

Witz S, Jung B, Fürst S, Möhlmann T (2012) De novo pyrimidine nucleotide synthesis mainly occurs outside of plastids, but a previously undiscovered nucleobase importer provides substrates for the essential salvage pathway in Arabidopsis. Plant Cell 24: 1549–1559

Yano R, Nakamura M, Yoneyama T, Nishida I (2005) Starch-related α-glucan/water dikinase is involved in the cold-induced development of freezing tolerance in Arabidopsis. Plant Physiol 138: 837–846

Zimmer D, Schneider K, Sommer F, Schroda M, Mühlhaus T (2018) Artificial intelligence understands peptide observability and assists with absolute protein quantification. Front Plant Sci 9:1559. doi: 10.3389/fpls.2018.01559

